# NLRP3 inflammasome priming and activation are regulated by a novel phosphatidylinositol-dependent mechanism

**DOI:** 10.1101/2020.01.14.905075

**Authors:** Claire Hamilton, Antoni Olona, Stuart Leishman, Kelly MacDonald-Ramsahai, Shamshad Cockcroft, Gerald Larrouy-Maumus, Paras K. Anand

## Abstract

Imbalance in lipid homeostasis is associated with discrepancies in immune signalling and is tightly linked to metabolic disorders. The diverse ways in which lipids impact immune signalling, however, remain ambiguous. The phospholipid phosphatidylinositol (PI), which is implicated in numerous immune disorders, is chiefly defined by its phosphorylation status. By contrast, the significance of the two fatty acid chains attached to the PI remains unknown. Here, by employing a mass-spectrometry-based assay, we demonstrate a role for PI acyl group chains in regulating both the priming and activation steps of the NLRP3 inflammasome in mouse macrophages. In response to NLRP3 stimuli, cells deficient in ABC transporter ABCB1, which effluxes lipid derivatives, revealed defective inflammasome activation. Mechanistically, *Abcb1*-deficiency shifted the total PI configuration exhibiting a reduced ratio of short-chain to long-chain PI-acyl lipids. Consequently, *Abcb1*-deficiency resulted in rapid degradation of TIRAP, the TLR adaptor protein which binds PI(4,5)-phosphate. Moreover, this accompanied increased NLRP3 phosphorylation at the Ser293 position and blunted inflammasome activation. Exogenously supplementing WT cells with linoleic acid, but not arachidonic acid, reconfigured PI acyl chains. Accordingly, linoleic acid supplementation increased TIRAP degradation, elevated NLRP3 phosphorylation, and abrogated inflammasome activation. Altogether, our study reveals a novel metabolic-inflammatory circuit which contributes to calibrating immune responses.

## Introduction

The NLRP3 inflammasome is a multiprotein complex activated in response to diverse pathogen-associated and ‘danger’ signals, and plays important roles during infectious and inflammatory diseases (Lupfer and Anand, 2016; Man and Kanneganti, 2015). The activation of NLRP3 inflammasome involves two steps in which NLRP3 is first licensed downstream of TLR signalling. This step, also known as the priming step, has both NF-κB-dependent and - independent consequences during NLRP3 activation. Upon sensing an apt stimulus, NLRP3 complexes with pro-caspase-1 and the adaptor molecule ASC. Consequently, caspase-1 is activated by auto-proteolysis, which further results in the maturation and release of biologically active forms of cytokines IL-1β and IL-18. Additional regulation is mediated by the post-translational modification of distinct NLRP3 domains which may affect either the priming or activation of the NLRP3 inflammasome (Lopez-Castejon et al., 2013; Yang et al., 2017). Inflammasome activation also results in the induction of inflammatory form of cell death, pyroptosis, by a Gasdermin-D-dependent mechanism (Kayagaki et al., 2015). Recent studies have revealed elaborate links between lipid metabolism and inflammasome activation (Anand, 2020). We, and others, previously demonstrated vital roles for cholesterol biosynthesis and transport in NLRP3 inflammasome activation (de la Roche et al., 2018; Guo et al., 2018). Other recent studies have demonstrated that NLRP3 recruitment to the dispersed *trans*-Golgi network requires binding to phosphoinositide 4-phosphate (PI4P) prior to NLRP3 activation (Chen and Chen, 2018). The mechanisms of NLRP3 inflammasome activation and, remarkably, the roles lipids play in the process remain ambiguous.

Lipid homeostasis is critical to all physiological processes including immune signalling (Barnett and Kagan, 2020). Lipids form the structural framework which imparts fluidity to membranes. Driven by their amphipathic nature, membrane lipids enable compartmentalization of cellular constituents both from the outside environment, and into discrete organelles (Casares et al., 2019). The lipid composition of membranes, in addition, is pivotal in shaping the localization, conformation, and thus the activity of lipid-protein and protein-protein complexes (Fernandis and Wenk, 2007). The latter is fundamental to cellular signalling emanating from the cholesterol-rich membrane-microdomains which serve as signalling platforms and are the preferred sites for pathogen entry (Kaul et al., 2004). Additionally, phosphoinositides (PIPs), the phosphorylated derivatives of the parent phosphatidylinositol (PI), play important roles in immune signalling by ensuring precise recruitment of the adaptor protein TIRAP, also known as Mal, to the activated TLRs at the plasma and endosomal membranes (Kagan and Medzhitov, 2006). Structurally, PI consists of an inositol head group and two fatty acyl chains linked by a glycerol backbone (Goud et al., 2016). The binding of PIPs to the phosphoinositide-binding domain of TIRAP results in the recruitment of MyD88 and the IRAK family of kinases to activated TLRs thereby promoting downstream NF-kB activation. Accordingly, the synthesis, turnover, and localization of PIPs may influence TLR-dependent signalling.

Homeostasis of the cellular lipid composition is largely maintained by the SREBP family of transcription factors that transcribe genes involved in lipid uptake, biosynthesis, and efflux (Horton et al., 2002). When in excess, lipids are either stored in the form of lipid droplets or are effluxed by the activity of the ABC family of transporters. The family members ABCA1 and ABCG1 facilitate cholesterol efflux to apolipoproteins and HDL, respectively (Yvan-Charvet et al., 2010). Moreover, deficiency in ABCA1 and ABCG1 is associated with enhanced secretion of inflammatory mediators, including IL-1β (Westerterp et al., 2017). Correspondingly, the efflux transporters play anti-inflammatory functions in diverse diseases (He et al., 2020; Sonett et al., 2018) suggesting a possible link between lipid metabolism and immune signalling (Castrillo et al., 2003). Cells deficient in *Abca1* and *Abcg1* cannot unload surplus lipids exhibiting elevated cholesterol accumulation which, independently, improves cytokine secretion (Westerterp et al., 2017; Yvan-Charvet et al., 2010). Overall, this argues that immune signalling is profoundly determined by lipid metabolism, but cholesterol accumulation remains a confounding factor and detailed mechanisms remain poorly defined. In this study, we investigated the role of lipid metabolism in immune signalling by studying ABCB1. ABCB1 (or P-glycoprotein) is a well-characterized family member that imparts multi-drug resistance to malignant cells but has no impact on cellular cholesterol levels (To et al., 2014; Vaidyanathan et al., 2016). The precise functions of ABCB1 in inflammasome activation remain undefined.

Here, we demonstrate a role for ABCB1 and PI fatty acyl chains in regulating the NLRP3 inflammasome. Mechanistically, *Abcb1*^-/-^ cells displayed a reduced ratio of short-chain to long-chain PI acyl chain lipids. This change in PI configuration was independent of the expression of enzymes that both synthesize PI and are involved in acyl chain remodelling. Remarkably, the shift in acyl chain composition regulated both NLRP3 priming and activation steps: it resulted in the depletion of TLR adaptor protein, TIRAP, and additionally elevated phosphorylation in the NACHT domain of NLRP3. Intriguingly, exogenously supplementing WT cells with linoleic acid reconfigured PI acyl chains, which accordingly mimicked *Abcb1*-deficient cells in TIRAP depletion, NLRP3 phosphorylation, and blunted inflammasome activity. Our study thus identifies an important role for PI lipid chain configuration in modulating inflammasome activity which may have significant implications in metabolic diseases.

## Results

### ABC transporter B family member 1 is required for caspase-1 activation and IL-1β secretion

Lipids are involved in diverse functions including maintenance of membrane structure, cellular signalling, and immunity (Barnett and Kagan, 2020; Casares et al., 2019). Consequently, altered lipid levels can lead to metabolic and inflammatory disorders, or may result in lipotoxicity (Sonett et al., 2018). Therefore, lipid homeostasis, and in particular, lipid efflux, is critical to normal cell function. Members of the ABC family function to export substrates, mainly lipids and related molecules, out of the cytosol (Yvan-Charvet et al., 2010). For example, ABCA1 is primarily associated with phospholipid and cholesterol transport to lipid-poor ApoA-I, while ABCG1 exports cholesterol to more mature HDL particles. Remarkably, deficiency in *Abca1* and *Abcg1* results in elevated secretion of inflammatory mediators in response to pathogenic stimuli (Westerterp et al., 2017). To address the roles of lipid metabolism in immune signalling, we investigated the role of ABCB1 in inflammasome activation. ABCB1 alters lipid metabolism and has a broad substrate specificity in transporting a range of drugs (Romsicki and Sharom, 2001). However, the transporter does not affect overall cellular lipid or cholesterol levels.

In order to test the role ABCB1 plays in inflammasome activation, we generated genetic knock-out cell lines using the CRISPR/Cas9 approach in immortalised bone marrow-derived macrophages (iBMDMs). In contrast to humans where a single isoform of ABCB1 is expressed, the mouse genome expresses two isoforms, *Abcb1a* and *Abcb1b* (Kalabis et al., 2005). We first examined the expression of the two isoforms in bone-marrow-derived mouse macrophages. PCR amplification followed by gel electrophoresis revealed that both *Abcb1a* and *Abcb1b* are expressed in unstimulated and LPS-stimulated iBMDMs **(Fig. S1A and S1B)**. However, quantitative PCR revealed that *Abcb1b* is expressed at approximately 600-fold higher levels than that of *Abcb1a*, suggesting that this isoform predominates in mouse macrophages **(Fig S1C)**. Similar results were obtained with primary mouse macrophages **(Fig S1D** and data not shown). Additionally, the expression of *Abcb1b* was only marginally higher in LPS-stimulated macrophages compared to control cells at all time-points tested, though there was a slight initial decrease in expression upon LPS stimulation **(Fig S1E)**. These studies, therefore, suggest a potential role for ABCB1 in immune cells.

Two cell lines, *Abcb1b*^-/-^ #1 and #2, were established with deletions in exon 10 of the protein, as verified by Sanger sequencing **(Fig S1F)**. The mRNA expression of *Abcb1b* in the two CRISPR cell lines was reduced significantly though the PCR amplicon was still synthesized to some extent **(Fig S1G)**. However, compared to WT cells, the protein expression of ABCB1 was diminished by more than 70% and 90% in the two cell lines potentially suggesting the presence of unstable mRNA in edited cell lines **(Figs 1A and 1B)**.

**Fig 1.**
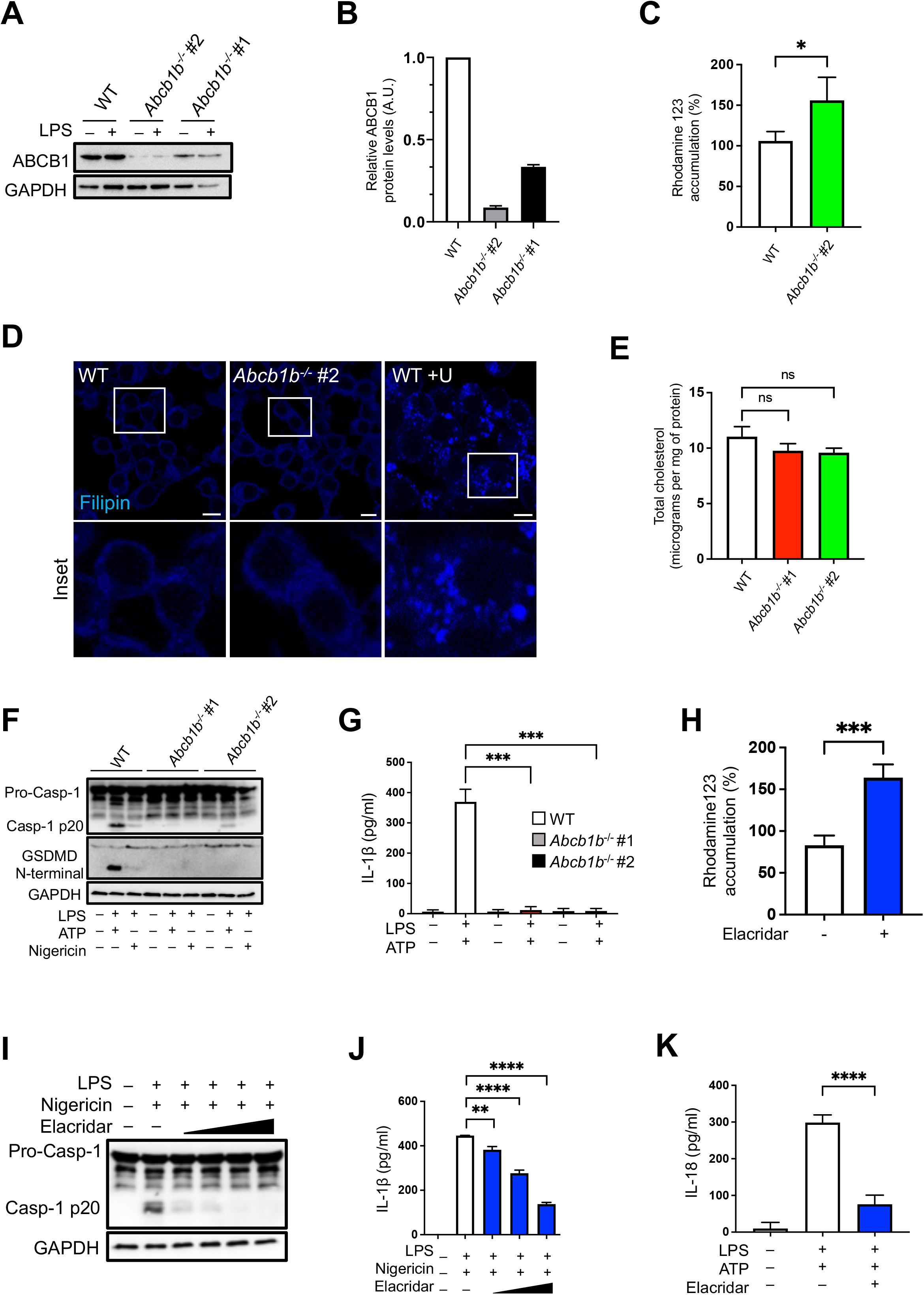
ABC transporter B family member 1 is required for caspase-1 activation and IL-1β secretion. **(A)** Cell lysates from WT, *Abcb1b*^-/-^ #1 and #2 cells were treated with or without LPS (500 ng/ml) for 4 hours and immunoblotted for ABCB1 and GAPDH. **(B)** Relative ABCB1 protein levels in WT, *Abcb1b*^-/-^ #1 and #2 cells measured with Image J on blots shown in (A). **(C)** WT and *Abcb1b*^-/-^ #1 and #2 cells were incubated with Rhodamine 123 (1 μM) for 30 minutes before measuring Rho123 accumulation on a fluorescent plate reader. **(D)** WT cells without and with overnight exposure to U18666a (5 μg/ml) and control *Abcb1b*^-/-^ cells grown on coverslips were fixed in paraformaldehyde for 1 hour followed by overnight staining with filipin (25 μg/ml) at 4 °C. Images were taken by confocal microscopy. **(E)** Total cholesterol content in WT, *Abcb1b*^-/-^ #1 and #2 cells measured by Amplex red assay. **(F)** WT, *Abcb1b*^-/-^ #1 and *Abcb1b*^-/-^ #2 cells were primed with LPS (500 ng/ml) for 4 hours followed by treatment with either ATP (5 mM) or nigericin (20 nM) for approximately 45 minutes. Cell lysates were collected and immunoblotted for caspase-1, GSDMD, and GAPDH. **(G)** Supernatants from macrophages treated as in (F) were analysed for IL-1β secretion by ELISA. **(H)** Primary mouse bone marrow-derived macrophages (BMDMs) were exposed to elacridar overnight (10 μM) and incubated with Rhodamine 123 (1 μM) for 30 minutes to evaluate Rh123 accumulation as in (C) above. **(I)** BMDMs were treated with increasing amounts of elacridar overnight (1, 2, 5 and 10 μM), followed by LPS priming (500 ng/ml) for 4 hours and nigericin (20 nM) for approximately 45 minutes. Cell lysates were immunoblotted for caspase-1 and GAPDH. **(J)** Supernatants from cells treated as in (I) were analysed for IL-1β secretion by ELISA. **(K)** Immortalised BMDMs (iBMDMs) were treated with elacridar overnight (10 μM) followed by LPS priming (500 ng/ml) for 4 hours and ATP (5 mM) for approximately 45 minutes. Cell supernatants were analysed for IL-18 secretion by ELISA. Data shown are mean ± SD, and the experiments shown are representative of at least three independent experiments. *, p = <0.05; **, p = <0.01; ***, p = <0,001, ****, p = <0.0001, by Student’s t-test.

To test the activity of ABCB1 in edited cells, we made use of Rhodamine 123 (Rho123) dye. Rho123 passively diffuses across biological membranes but is subsequently metabolised by intracellular esterases to yield a fluorescent compound that has reduced permeability (Forster et al., 2012). The efflux of the dye out of cells requires active transport by ABCB1 and certain other ABC transporters (Forster et al., 2012). *Abcb1*^-/-^ cells exposed for 30 min to Rho123 showed increased sequestration of the dye compared to control cells thereby confirming inhibition of ABCB1 transporter activity **(Fig 1C)**.

Unlike other ABC transporters that are involved in lipid efflux, ABCB1 is not known to alter cellular cholesterol levels (Garrigues et al., 2002). In agreement, WT and *Abcb1b*^-/-^ macrophages exhibited no qualitative differences in cholesterol esterification upon staining with filipin, a compound that binds to free unesterified cholesterol (de la Roche et al., 2018). Excitation of filipin by UV fluorescence showed no obvious differences in either the levels or distribution of cholesterol between WT and *Abcb1b*^-/-^ cells **(Fig 1D)**. As a control, we exposed WT cells to U18666a, which specifically blocks the lysosomal cholesterol transporter, NPC1 (de la Roche et al., 2018), and this resulted in expected punctate filipin staining suggestive of lysosomal cholesterol accumulation **(Fig 1D)**. Additionally, quantitative analysis of total cellular cholesterol revealed similar levels in both WT and *Abcb1b*^-/-^ cells **(Fig 1E)**. These data agree with previous studies and indicate no major effect of ABCB1 deficiency on cholesterol esterification, distribution, and efflux.

We next tested the activation of NLRP3 inflammasome in macrophages with genetic deletion of *Abcb1b*. WT and *Abcb1*^-/-^ cells were exposed to LPS for 4h followed by either ATP or nigericin. NLRP3 inflammasome activation results in the assembly of the inflammasome complex containing NLRP3, the adaptor ASC, and pro-caspase-1. Pro-caspase-1 is then cleaved into its active p20 form, which subsequently results in the maturation and secretion of pro-inflammatory cytokines, IL-1β and IL-18. Compared to WT cells, *Abcb1b*^-/-^ cells exhibited diminished caspase-1 cleavage and IL-1β secretion upon exposure to both nigericin and ATP **(Figs 1F and 1G)**. Activation of the NLRP3 inflammasome also leads to an inflammatory form of cell death, termed pyroptosis, which is mediated by gasdermin-D (GSDMD). Cleavage of GSDMD by caspase-1 results in an N-terminal fragment, which subsequently polymerises and forms pores in the cell membrane, ultimately leading to cell rupture. GSDMD cleavage in *Abcb1b*^-/-^ macrophages was diminished in comparison to WT cells following NLRP3 inflammasome activation, further corroborating reduced caspase-1 in deficient cells **(Fig 1F)**.

We next used elacridar, a third-generation inhibitor, that has been shown to inhibit ABCB1 activity and overcome drug resistance in cancer models (Dash et al., 2017). Primary BMDMs treated overnight with increasing concentrations of elacridar and exposed for 30 min to Rho123 exhibited increased sequestration of the dye compared to control cells **(Fig 1H)**. In agreement with deficient cells, treatment of WT macrophages with increasing concentrations of elacridar followed by NLRP3 activation resulted in a dose-dependent decrease in the generation of cleaved caspase-1 p20 form **(Fig 1I)**, and reduction in IL-1β and IL-18 secretion **(Figs 1J and 1K)**. Similar results were observed with the P2X_7_ receptor agonist and NLRP3 activator, ATP **(Figs S2A and S2B, and Fig 1K)**. Together, these data demonstrate the functional requirement of ABCB1 in NLRP3 inflammasome activation.

### ABCB1 is required for the priming and activation of the NLRP3 inflammasome

NLRP3 inflammasome activation requires two distinct steps. The priming step results in the induction of TLR-dependent NF-κB activation, which in turn leads to the transcriptional and translational upregulation of NLRP3 and pro-IL-1β. The second signal activates the NLRP3 inflammasome complex, leading to caspase-1 activation (Hamilton et al., 2017; Hamilton and Anand, 2019). We next studied the precise step at which ABCB1 is required for activation of the NLRP3 inflammasome by examining the expression of inflammasome components including NLRP3, and pro-IL-1β. Stimulation with LPS and ATP upregulated the NLRP3 and pro-IL-1β protein in WT cells, which was found to be blunted in cells lacking *Abcb1* **(Fig 2A)**. By contrast, pro-caspase-1 and ASC are constitutively expressed, and their expression remained unaltered in *Abcb1b*^-/-^ macrophages **(Figs 1F and 2A)**. These results coincided with reduced upregulation of *Nlrp3* and *Il1b* at the mRNA level **(Figs 2B and 2C)**. NF-κB is inactive within the cell prior to stimulation through binding to the inhibitory protein IkB. Upon TLR stimulation, signal transduction leads to the activation of the IKK complex, which subsequently phosphorylates IkB, targeting it for degradation. This allows NF-κB to translocate to the nucleus and initiate gene transcription. In addition to NF-κB activation, TLR stimulation also leads to the activation of MAPKs, notably p38 (Kawasaki and Kawai, 2014). In agreement with the above data, the phosphorylation of IkB and p38 MAPK was reduced in LPS-stimulated *Abcb1b*^-/-^ cells compared to WT cells **(Fig 2D)**. We also examined the expression of other NF-κB -dependent cytokines and chemokines in response to TLR4 activation and their expression were similarly decreased in cells lacking ABCB1 **(Figs 2E-G)**. Moreover, the mRNA expression of interferon-β, which is dependent on adaptor TRIF upon LPS stimulation, was also reduced in *Abcb1b*^-/-^ cells **(Fig 2H)**. Furthermore, this response was not specific to TLR4 ligation, as activation of TLR2 or TLR7 by Pam3CSK4 and imiquimod, respectively also resulted in diminished upregulation of *Nlrp3, Il1β, Tnfα* and *Cxcl1* (KC) mRNA expression **(Fig 2I, and Figs S3A-C)**. These data suggested blunted TLR-dependent signalling in the absence of ABCB1.

**Fig 2.**
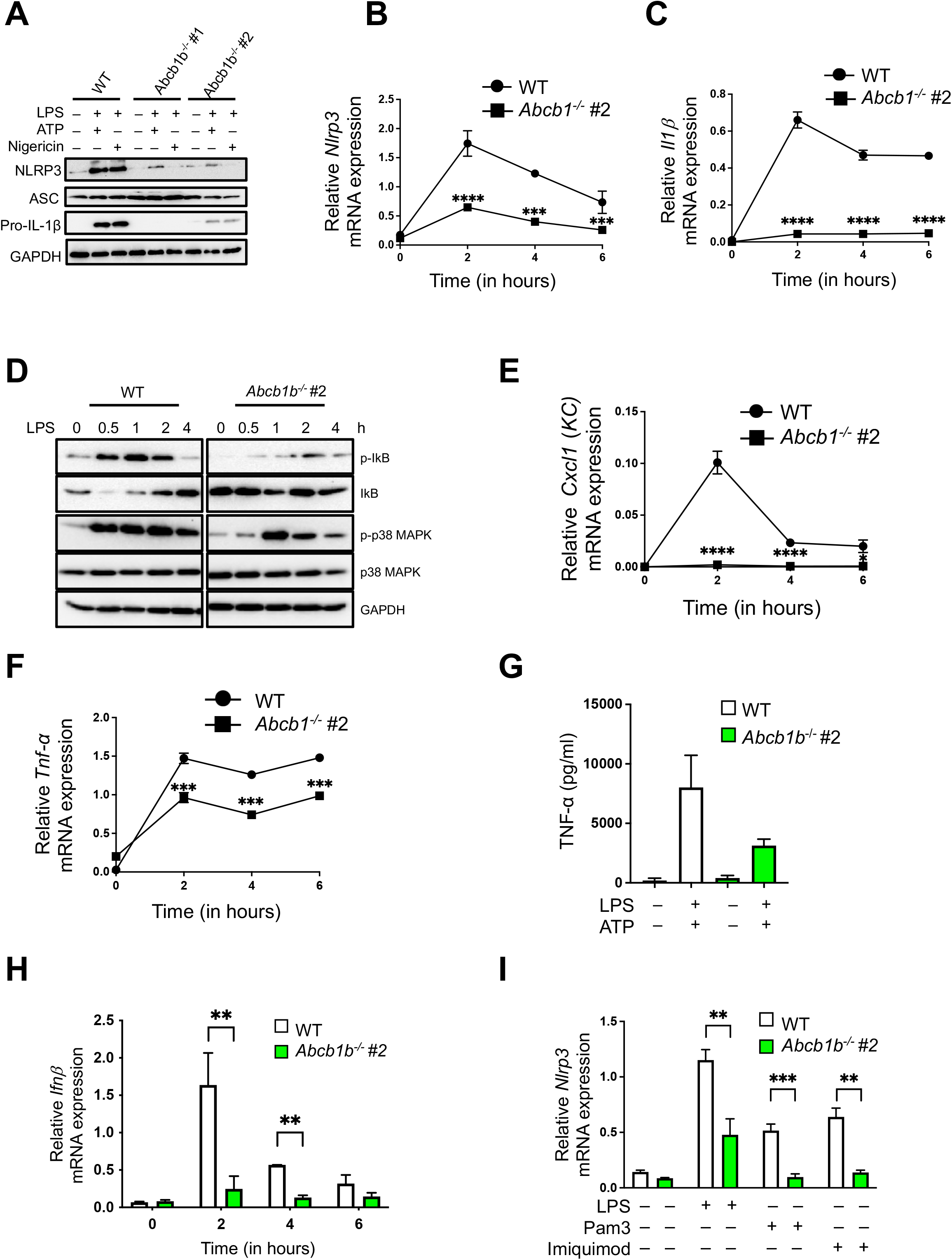
ABCB1 is required for the priming and activation of the NLRP3 inflammasome. **(A)** WT, *Abcb1b*^-/-^ #1 and #2 cells primed with LPS (500 ng/ml) for 4 hours followed by treatment with either ATP (5 mM) or nigericin (20 nM) for approximately 45 minutes. Cell lysates were collected and immunoblotted for NLRP3, ASC, pro-IL-1β and GAPDH. **(B)** WT and *Abcb1b*^-/-^ #2 cells were stimulated with LPS (500 ng/ml) for the indicated time points. RNA was extracted and converted into cDNA. Gene expression of *Nlrp3* and **(C)** *Il1β* was determined by real-time PCR. **(D)** WT and *Abcb1b*^-/-^ #2 cells were stimulated with LPS (500 ng/ml) for the indicated time points. Protein samples were collected and immunoblotted for phospho-IκB, total IκB, phospho-p38 MAPK, total p38 MAPK and GAPDH. **(E)** Cells treated as in B were examined for *Cxcl1 (KC*) and **(F)** *Tnfα* gene expression by real-time PCR. **(G)** Supernatants from WT and *Abcb1b*^-/-^ #2 macrophages treated as in (B) were analysed for TNF-α cytokine secretion by ELISA. **(H)** WT and *Abcb1b*^-/-^ #2 cells were stimulated with LPS (500 ng/ml) for 4 hours and analysed for mRNA expression of *Ifnβ*. mRNA expression is shown relative to *Gapdh*. **(I)** WT and *Abcb1b*^-/-^ #2 cells were stimulated with either LPS (500 ng/ml), Pam3 (1 μg/ml) or Imiquimod (1 μg/ml) for 4 hours and analysed for *Nlrp3* mRNA expression. Gene expression is shown relative to *Gapdh*. Data shown are mean ± SD, and the experiments shown are representative of 3 independent experiments. *, p = <0.05; **, p = <0.01; ***, p = <0.001; ****, p = <0.0001, by Student’s t-test.

### ABCB1 does not affect the activation of NLRC4 and AIM2 inflammasomes

We next tested the requirement of ABCB1 during NLRC4 and AIM2 inflammasome activation. Unlike NLRP3, the expression of NLRC4 and AIM2 is constitutively expressed at significantly higher levels in mouse macrophages. However, the expression and therefore the secretion of downstream effector cytokine IL-1β still requires upregulation by TLR signalling. Activation of NLRC4 inflammasome by *Salmonella* infection and AIM2 inflammasome by exposure to poly(dA:dT) resulted in comparable caspase-1 cleavage between WT and *Abcb1b*^-/-^ cells **(Fig 3A** and data not shown). However, the levels of IL-1β secretion, as expected, were found diminished in both *Abcb1b*^-/-^ cells and in WT cells where ABCB1 was pharmacologically blocked during NLRC4 and AIM2 inflammasome activation **(Figs 3B and 3C)**. In contrast to pro-IL-1β, pro-IL-18 is constitutively expressed and does not require TLR signalling for upregulation. In agreement, no difference in IL-18 production was observed following NLRC4 or AIM2 inflammasome activation in cells lacking ABCB1 **(Fig 3D)**. Secretion of IL-18, however, following NLRP3 activation showed complete abolishment in *Abcb1b*^-/-^ macrophages **(Fig 3D)**. These data suggest that the function of ABCB1 is specific to the NLRP3 inflammasome, and that the diminished IL-1β production is due to dampened NF-κB signalling in *Abcb1b*^-/-^ cells **(Figs 2C and 2D)**.

**Fig 3.**
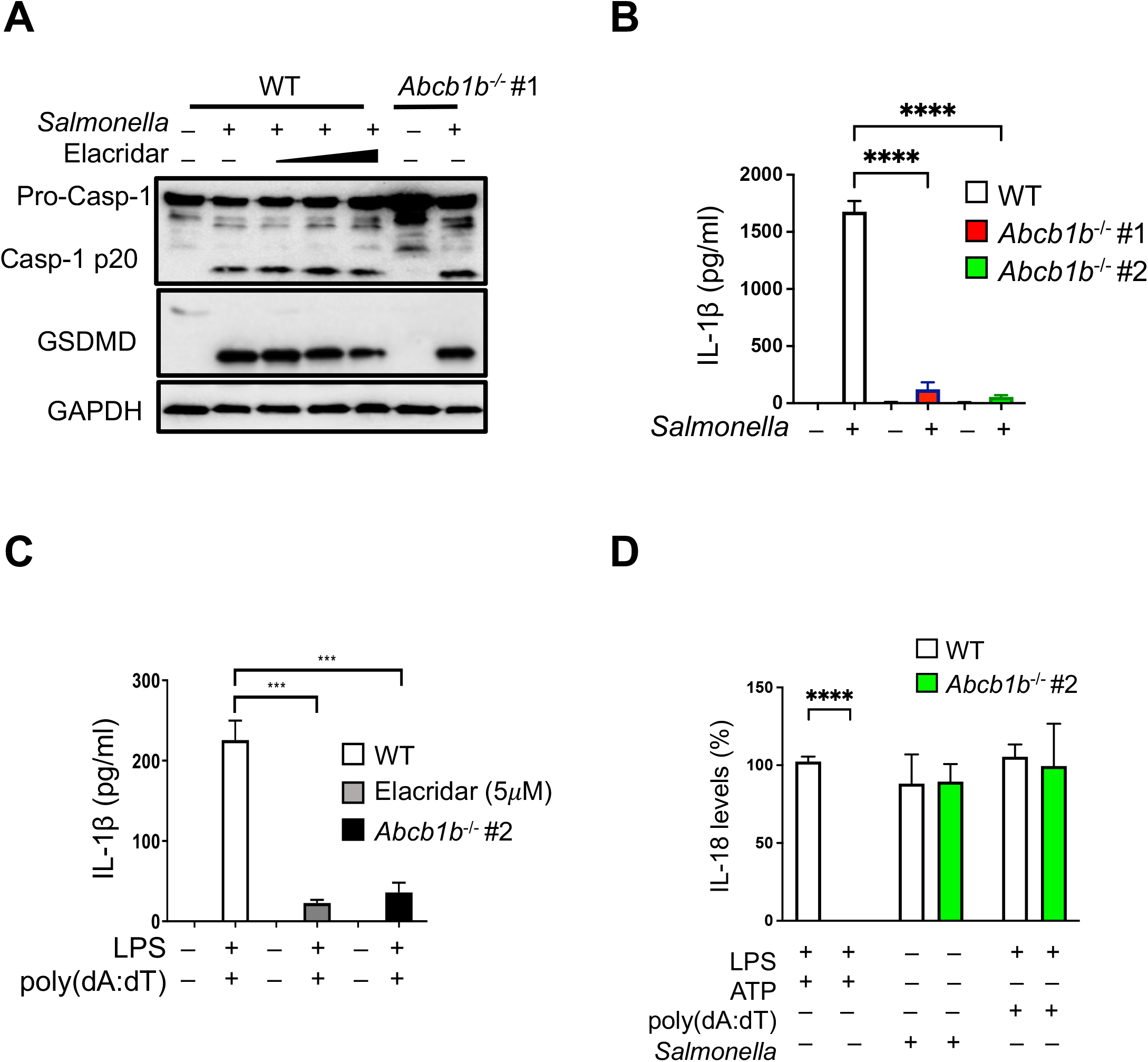
ABCB1 does not affect the activation of NLRC4 and AIM2 inflammasomes. **(A)** WT cells with or without exposure to elacridar (2, 5, 10 μM; 16 hours) and *Abcb1b*^-/-^ #1 cells were infected with *Salmonella typhimurium* at an MOI of 2 for approximately 4-5 hours. Cell lysates were collected and immunoblotted for caspase-1, GSDMD and GAPDH. **(B)** WT, *Abcb1b^-/-^ #1*, and *Abcb1b*^-/-^ #2 macrophages were infected with *Salmonella typhimurium* at an MOI of 2 for approximately 4-5 hours and supernatants were analysed for IL-1β by ELISA. **(C)** Cells were treated with LPS (500 ng/ml) for 4 hours followed by transfection with 1 μg of DNA complexed to Lipofectamine 2000 (ratio DNA:Lipofectamine2000: 1:3) for approximately 4 hours to activate the AIM2 inflammasome. Supernatants were analysed for IL-1β by ELISA. **(D)** Supernatants from cells treated as in (B) and (C) or treated with LPS (500 ng/ml, 4 hours) and ATP (5 mM, 45 minutes) were analysed for IL-18 production by ELISA. Data shown are mean ± SD and is representative of at least three independent experiments. **, p=<0.01, *** p = <0.001, **** p = <0.0001, by Student’s t-test.

### Blunted NLRP3 inflammasome activation is associated with a shift in PI lipid chains

A notable feature of the plasma membrane is the distinct lipid composition in the two leaflets of the bilayer (Van Meer et al., 2008). Sphingolipids are mostly present in the outer leaflet while glycerophospholipids such as phosphatidylethanolamine (PE), phosphatidylserine (PS), and phosphatidylinositol (PI) are mostly restricted to the inner leaflet thereby imparting distinct functions to the membranes (Van Meer et al., 2008).

To further investigate the mechanism by which *Abcb1*-deficiency dampens NLRP3 activation, we performed whole-cell lipidomics by MALTI-TOF MS. Our experiments revealed ions in the 800 – 1700 *m/z* range. The most striking change was observed in the 800-920 *m/z* range which represents the total PI complement present in cells with distinct signals corresponding to anticipated PI masses. In mammalian cells, a large fraction of the PIP molecules has saturated stearic acid (18:0) at the *sn*-1 position and unsaturated arachidonic acid (20:4) at the *sn*-2 position **(Fig 4A)**. Consequently, both the WT and *Abcb1*^-/-^ cells exhibited a peak at *m/z* 885.4, described together as PI (38:4) **(Figs 4B and 4C)**. Additionally, another PI peak at *m/z* 861.4, corresponding to PI (36:2), was observed in WT and *Abcb1*^-/-^ cells **(Figs 4B and 4C)**. MS/MS fragmentation of the peaks at *m/z* 861 and *m/z* 885 revealed the presence of stearic acid, arachidonic acid, and oleic acid **(S4 Fig)**. Besides the presence of PIs with distinct acyl chain lengths, other closely related PI siblings were also represented though at much lower levels **(Figs 4B and 4C, and Fig S5)**. Notably, the WT cells exhibited almost equal levels of the two PI-lipid masses at 861 and 885. By contrast, *Abcb1*-deficient cells favoured higher levels of PI (38:4) compared to PI (36:2) which was expressed at significantly lower levels. As a result, the ratio of the two PI-lipid masses consistently demonstrated a significant difference between WT and *Abcb1*^-/-^ cells **(Fig 4D)**. These studies, therefore, suggest that the deficiency in *Abcb1* modifies the PI fatty acyl configuration resulting in a reduced ratio of short-chain to long-chain fatty acids. **(Fig 4D)**.

**Fig 4.**
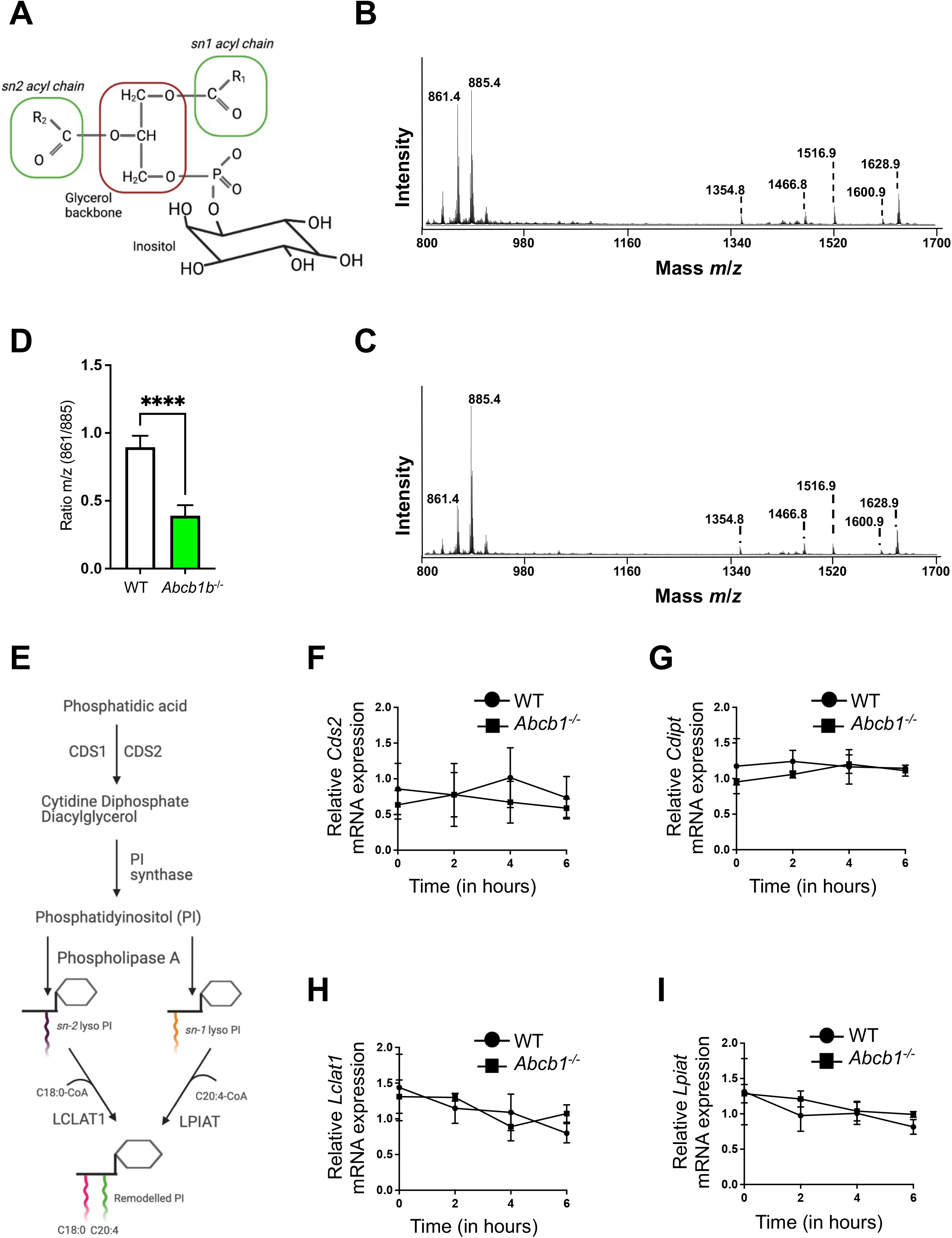
Blunted NLRP3 inflammasome activation is associated with a shift in PI lipid chains. **(A)** Phosphatidylinositol structure consists of an inositol headgroup, a glycerol backbone (red box), and two acyl chains R_1_ and R_2_ at the *sn-1* and *sn-2* positions (green boxes). Representative spectra of **(B)** WT and **(C)** *Abcb1b*^-/-^ cells. Peaks of interest are indicated. The peaks at m/z 835.4, 861.4, 885.4 and 911.5 are assigned to phosphatidyl-inositol (PI) 34:1, 36:2, 38:4 and 40:4, respectively. In this range are also found peaks at m/z 1354.7, m/z 1438.8 and m/z 1466.8, which are assigned to GM-2 d18:1/16:0, GM-2 d18:1/22:0, and GM-2 d18:1/C24:0 respectively. In the range m/z 1500-1650 are found GM-1 at m/z 1516.8, m/z 1544.8, m/z 1572.8, m/z 1600.9, m/z 1626.9 and m/z 1628.9 assigned to GM-1 18:1/16:0, 18:1/18:0, 18:1/20:0, 18:1/22:0, 18:1/24:1 and 18:1/24:0, respectively. **(D)** Ratio m/z of peaks 861/885 in WT and *Abcb1b*^-/-^ macrophages. **(E)** Model showing PI synthesis, and subsequent remodelling to acquire typical fatty acid configuration. WT and *Abcb1b*^-/-^ #2 cells were treated with LPS (500 ng/ml) for the indicated time points, and the mRNA expression of **(F)** *Cds2*, **(G)** *Cdipt*, **(H)** *Lclat1*, and **(I)** *Lpiat* were analysed. Gene expression is shown relative to *Gapdh*. Data shown are mean +/- SD and is representative of at least three independent experiments. **, p=<0.01, *** p = <0.001, **** p = <0.0001, by Student’s t-test.

Further analysis of the mass-spec data also unveiled several ganglioside peaks that were distinct between WT and *Abcb1b*^-/-^ macrophages. In particular, the percentage of GM1 ganglioside was significantly reduced in *Abcb1b*^-/-^ cells **(Fig S6A)** which was further confirmed qualitatively and quantitatively by labelling WT and *Abcb1*^-/-^ cells with cholera toxin subunit B (CTB) or by flow cytometry **(Figs S6B-F)**.

The *de novo* synthesis of PI occurs at the ER through the conversion of phosphatidic acid (PA) via 2 enzymatic reactions **(Fig 4E)**. PA is first converted into an intermediate, cytidine diphosphate-diacylglycerol (CDP-DAG), by CDP-DAG synthase (CDS). CDP-DAG is then converted to PI by PI-synthase (also known as CDIPT) (Blunsom and Cockcroft, 2020; Le Guédard et al., 2009). Cleavage of the fatty acid by phospholipases A, followed by acylation by acyl-CoA–specific lysophospholipid acyltransferase enzymes, is needed to incorporate a new fatty acid molecule (Barneda et al., 2019). Lysophosphatidylinositol acyltransferase (LPIAT) has been shown to reacylate PI with polyunsaturated substrates such as arachidonic acid, while lysocardiolipin acyltransferase 1 (LCLAT1) has been demonstrated to reacylate with stearic acid (Barneda et al., 2019) **(Fig 4E)**. As *Abcb1b*-deficient macrophages displayed altered PI lipid chains, we investigated whether this was due to a defect in PI synthesis and or acyl chain remodelling. However, the mRNA expression of *Cds2* (*Cds1* was expressed with Ct value >30) and *Cdipt* revealed no significant difference between WT and *Abcb1b*^-/-^ cells **(Figs 4F and 4G)**. Similarly, the mRNA expression of *Lclat1* and *Lpiat* remained unchanged in deficient cells **(Figs 4H and 4I)**. These data indicate that ablation of *Abcb1* in macrophages alters PI lipid chain configuration independently of the synthesis of PI or remodelling of its acyl chain composition.

### Exogenous supplementation with linoleic acid alters PI acyl chain configuration

To further understand the mechanistic underpinnings of our findings and to validate them, we next sought to identify the fatty acids which may alter PI acyl chains and subsequently modify inflammasome activity. In agreement with previous studies (Hunt et al., 2004), we hypothesized that exogenous supplementation with distinct fatty acids will alter cellular PI acyl chain conformation. To achieve this, cells were grown for at least two weeks in growth media supplemented with different fatty acids prior to validating the samples by mass spectrometry **(Fig 5A)**. These assays revealed that prolonged supplementation with linoleic acid, but not arachidonic acid, favoured long-chain fatty acids **(Fig 5A)**. This was predominantly reflected in the increased concentration of unsaturated acyl chains at the *sn*-2 position. Compared to control cells, supplementation with linoleic acid changed the fatty acid composition to that observed in *Abcb1*^-/-^ cells **(Figs 5A, S4, and S5)**. Consequently, these cells behaved similarly to cells lacking ABCB1. Strikingly, supplementation with arachidonic acid did not result in any noticeable difference **(Fig S5)**. Notably, we also observed a narrower range over which macrophages tolerated arachidonic acid (data not shown). Accordingly, compared to control cells, WT cells grown in the presence of linoleic acid demonstrated reduced *Nlrp3* and *Il-1β* mRNA expression **(Figs 5B and 5C)**. Again, these results were specific for linoleic acid as supplementation with arachidonic acid, alone or in combination with stearic acid, neither altered the *m/z* 861/885 ratio nor the mRNA expression of *Nlrp3* and *Il-1β* **(Figs 5A, 5D and 5E, and S5)**. Furthermore, like *Abcb1*^-/-^ cells, exogenous linoleic acid resulted in the disruption of GM1 presence in WT cells **(Fig 5F and 5G)**. These results, therefore, demonstrate that linoleic acid supplementation alters the PI acyl chain profile.

**Fig 5.**
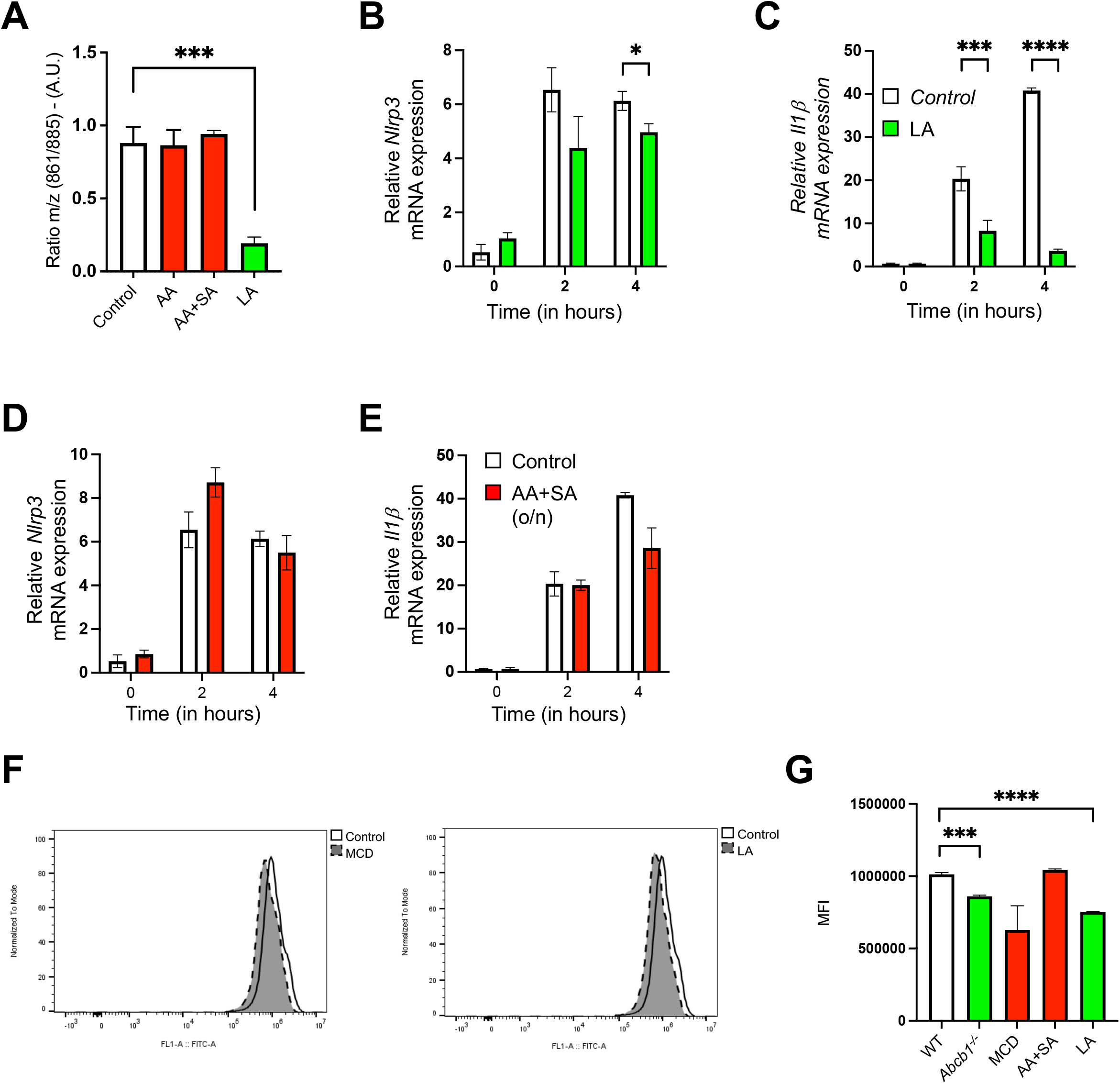
Exogenous supplementation with linoleic acid alters PI acyl chain configuration. WT and *Abcb1*^-/-^ cells were cultured in the presence of either arachidonic acid (5 μM, AA) or AA (5 μM) and stearic acid (20 μM, SA) or linoleic acid (20 μM, LA) for at least two weeks with the regular splitting of cell cultures every 2-3 days before subjecting the samples to whole-cell lipidomics. **(A)** Ratio *m/z* of peaks 861/885 in cells grown in the presence of indicated fatty acids. **(B-E)** WT cells cultured with the shown fatty acids were treated with LPS (500 ng/ml) for the indicated time points, and the mRNA expression of **(B, D)** *Nlrp3*, and **(C, E)** *Il-1β* were analysed. Gene expression is shown relative to *Gapdh*. **(F)** WT (and *Abcb1^-/-^*) cells either untreated or treated with MβCD (10 μm, 30 min) or the indicated fatty acids were stained with cholera toxin B (CTB) (1 μg/mL) for 10 minutes at 4 °C followed by incubation with Alexa Fluor 488-conjugated anti-CTB antibody for 15 minutes at 4 °C to reveal GM1 presence. Fluorescence was analysed by flow cytometry and representative spectra are shown. **(G)** MFI (Mean fluorescence intensity) quantification of cells treated as above. Data shown are mean ± SD and is representative of at least three independent experiments. *, p=<0.05, *** p = <0.001, **** p = <0.0001, by Student’s t-test.

### PI acyl chain profile regulates TIRAP expression

TLRs are transmembrane receptors that can sense microbial products at distinct subcellular sites. Ligation of TLRs at both the plasma membrane and endosomes triggers a signal transduction pathway involving the adaptor protein MyD88 which is recruited to the conserved TIR domain present in the cytosolic tails of these receptors. Most TLRs recruit MyD88 by employing the intermediate sorting adaptor TIRAP, which is bound to either PIP2 or PI4P for precise localization, and is prepositioned on the membranes prior to TLR activation. The phosphorylation by Bruton’s tyrosine kinase activates TIRAP but this is immediately followed by SOCS1-mediated TIRAP polyubiquitination and degradation to avoid sustained signalling (Mansell et al., 2006). The rapid turnover of TIRAP is similarly regulated by serine/threonine kinases, interleukin-1 receptor-associated kinases IRAK1 and IRAK4, which directly phosphorylate TIRAP triggering Lys48 -linked ubiquitination and proteasomal degradation (Dunne et al., 2010).

We next evaluated the role of altered PI-acyl chain composition on TLR signalling by assessing TIRAP expression. As reported in previous studies, we mostly found TIRAP at the cell periphery with additional punctate staining in the cytoplasm **(Fig. 6A)**. Stimulation of WT cells with LPS resulted in increased proximity of TIRAP to the plasma membrane **(Fig. 6A)**. LPS stimulation resulted in a modest increase in TIRAP expression as observed by WB at 15 min followed by sustained TIRAP expression up to 60 min, the last time point that we tested **(Fig. 6B and 6C)**. By contrast, cells lacking *Abcb1* or those exposed to linoleic acid, exhibited significant depletion in TIRAP expression **(Fig. 6B and 6C)**. Following phosphorylation, TIRAP is degraded by the 26S proteasome. Accordingly, pre-treatment of linoleic acid -supplemented cells with the proteasomal inhibitor MG132 restored TIRAP expression **(Fig. 6C)**. These studies, therefore, suggest that change in PI fatty acid configuration modifies TLR signalling by regulating the degradation of adaptor protein TIRAP.

**Fig 6.**
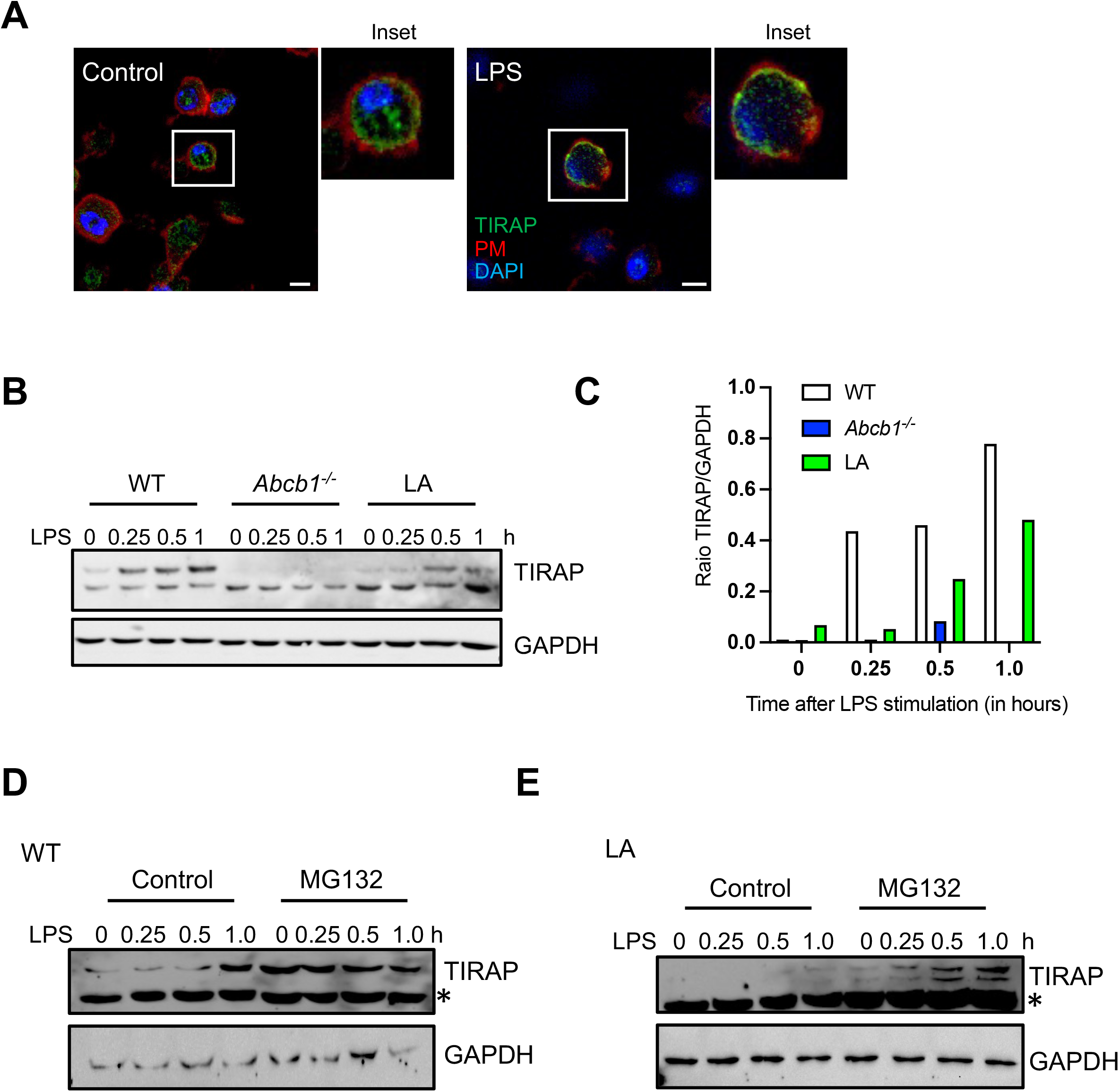
PI acyl chain profile regulates TIRAP expression. **(A)** Control and LPS-primed WT cells grown on coverslips were labelled with anti-TIRAP antibody and nuclei were stained with DAPI. The plasma membrane was stained by adding AF-647-conjugated phalloidin for the last 15 min. **(B)** WT, *Abcb1*^-/-^ and linoleic acid-supplemented cells were stimulated with LPS for different times. Cell lysates were collected and immunoblotted for TIRAP and GAPDH. **(C)** Quantitation of the WB shown in **(B)** by Image J. **(D)** WT and **(E)** linoleic acid-grown cells were pre-treated or not with the proteasomal inhibitor MG132 (10 μM) for 30 min before stimulating the cells with LPS for different time points. Cell lysates were collected and immunoblotted for TIRAP and GAPDH. The data shown are representative of at least three independent experiments. Asterisk (*) on immunoblots donates a non-specific band. Bars, 5 μm.

### Altered PI acyl chain profile blunts inflammasome activity by increasing NLRP3 phosphorylation

The NLRP3 inflammasome is regulated both at the transcriptional and post-translational levels. In particular, NLRP3 phosphorylation at distinct sites regulates both the priming and activation steps. Notably, linoleic acid derivative, prostaglandin E2, has been shown to facilitate NLRP3 phosphorylation in the NACHT domain at Ser293 position (Ser295 in humans) by inducing protein kinase A (Mortimer et al., 2016). Therefore, we next tested the status of NLRP3 Ser293 phosphorylation in control and linoleic acid -supplemented cells. We noted from different experiments that cells exposed to linoleic acid display on average 1.5 times less NLRP3 expression due to defective priming under these conditions. To ensure that we examined phosphorylation on equal NLRP3 protein levels, we increased the amount of protein loaded onto gels in linoleic acid -supplemented cell lysates. Accordingly, increasing the protein concentration by the above fraction resulted in similar total NLRP3 expression in both control and linoleic acid-supplemented samples **(Fig. 7A)**. By contrast, compared to control cells, linoleic acid-exposed cells demonstrated increased phosphorylation as examined using a phospho-NLRP3 specific antibody **(Fig. 7A and 7B)**. In agreement, linoleic acid -supplemented cells exhibited reduced caspase-1 activity **(Fig. 7C)**. However, exposure to arachidonic acid resulted in similar caspase-1 activity **(Fig. 7C)**. Moreover, linoleic acid supplementation diminished the secretion of IL-18 from LPS-primed cells in response to both ATP and nigericin **(Fig. 7D)**. In order to further validate our results, we next examined inflammasome assembly by examining ASC speck formation. NLRP3 inflammasome activation by LPS+ATP resulted in a comparable percentage of ASC specks in both control WT and cells exposed to arachidonic acid **(Fig. 7E and 7F)**. By comparison, cells lacking *Abcb1* exhibited significantly reduced ASC specks. Similarly, cells grown in linoleic acid-rich media displayed reduced ASC speck formation **(Fig. 7E and 7F)**. Together, these data demonstrate that altered PI acyl chain configuration affects both the priming and activation steps of the NLRP3 inflammasome.

**Fig 7.**
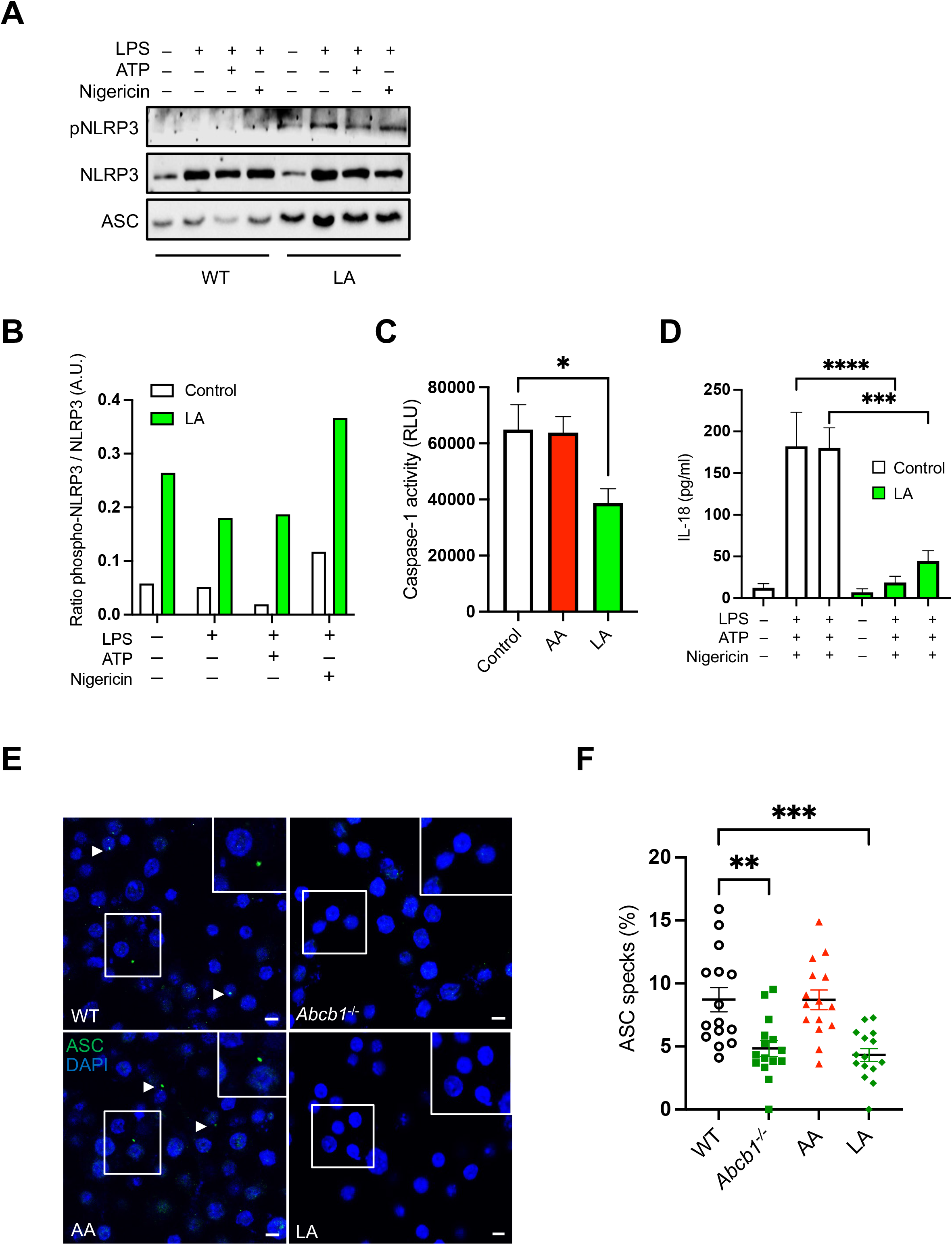
Altered PI acyl chain profile blunts inflammasome activity by increasing NLRP3 phosphorylation. **(A)** WT and linoleic acid-supplemented cells (LA) were either left untreated or primed with LPS (500 ng/ml) for 4h followed by treatment with either ATP (5 mM) or nigericin (20 nM) for approximately 45 minutes. Cell lysates were collected, and protein quantified. The amount of protein loaded for linoleic acid -supplemented cells was increased to normalize total NLRP3 levels. Cell lysates were immunoblotted for phosphor-NLRP3, total NLRP3, and ASC. **(B)** Quantitative analysis of the WB shown in **(A)** by Image J. **(C)** Control WT, arachidonic acid-grown cells (AA) and linoleic acid-grown cells (LA) were primed with LPS (500 ng/ml) for 4h followed by treatment with ATP (5 mM) for approximately 45 minutes. Caspase-Glo 1 activity was measured in the culture supernatants. **(D)** Cell supernatants collected as in **(A)** were assayed for IL-18 secretion by ELISA. **(E)** WT, *Abcb1*^-/-^, arachidonic acid-grown (AA) or linoleic acid-grown cells (LA) cells were exposed to LPS + ATP followed by labelling with anti-ASC antibody and DAPI staining. **(F)** Quantitative analysis of the percentage of cells with ASC specks in samples treated as above. Each dot represents an individual field with at least n = 40 cells. Data shown are mean ± SEM, and the experiments shown are representative of at least three independent experiments. Arrowheads show ASC specks. Bars, 5 μm. **, p=<0.01, *** p = <0.001, **** p, = <0.0001 by Student’s t-test.

## Discussion

Our studies described here demonstrate that lipid metabolism is intricately linked to immune signalling and inflammasome activation. Genetic or pharmacological depletion of ABCB1 abolished caspase-1 cleavage and IL-1β secretion. Remarkably, this was restricted to the NLRP3 inflammasome as *Abcb1*^-/-^ cells displayed comparable activation of the NLRC4 and AIM2 inflammasomes. Further mechanistic studies based on whole-cell lipidomics revealed an altered PI lipid chain profile in deficient cells. Remarkably, modified PI lipid chain configuration accompanied reduced TIRAP expression and altered NLRP3 phosphorylation; together they affected both NLRP3 inflammasome priming and activation steps **(Fig. 8)**. Notably, prolonged growth in media supplemented with linoleic acid reconfigured PI acyl chain profile in WT cells thereby mimicking the phenotypic features observed in deficient cells including aberrant inflammasome activity. Our results, therefore, demonstrate the metabolic regulation of inflammasome activation by PI acyl chains.

**Fig 8.**
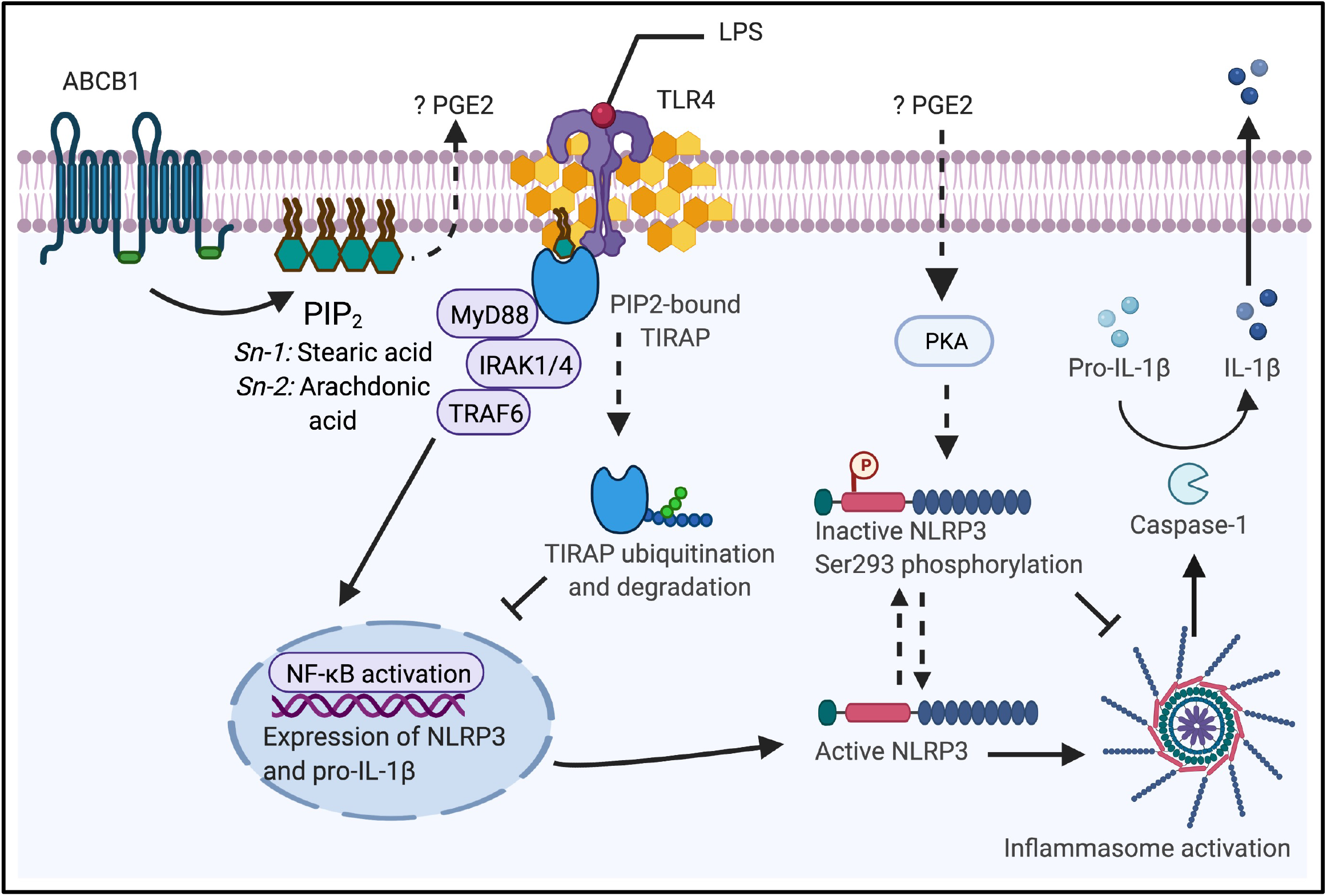
Schematic for ABCB1- and PI -mediated regulation of the NLRP3 inflammasome. ABCB1 is important in maintaining lipid metabolism. In the absence of ABCB1, the PI lipid chain configuration is altered resulting in the reduced ratio of short-chain to long-chain fatty acids. The activation of TLR4-dependent pathway relies on the adaptor protein TIRAP binding to PIP2 for precise positioning at the plasma membrane. However, alteration in PI-lipid profile results in at least two distinct outcomes which affect both the priming and activation steps of the NLRP3 inflammasome. First, either due to inability to bind to PIP2 and or reduced PIP2 abundance at the PM, TIRAP is ubiquitinated and degraded in the cytoplasm. Inflammasome assembly requires NLRP3, ASC, and pro-Caspase-1 in a complex wherein caspase-1 activation leads to the maturation of pro-IL-1β into the activation form. Altered PI profile most likely increases PGE2 secretion because of increased PI incorporation of precursor arachidonic acid. As a result, NLRP3 is phosphorylated at Ser293 residue which is mediated by protein kinase A (PKA) leading to NLRP3 inactivation. Consequently, assembly of the inflammasome is abrogated resulting in blunted caspase-1 activation and IL-1β secretion. PGE2, prostaglandin E2; TIRAP, Toll-interleukin-1 receptor (TIR) domain-containing adaptor protein; PIP2, phosphatidylinositol (4,5) bisphosphate.

PIPs, which take key roles in membrane trafficking and signal transduction, are defined by the phosphorylation of their inositol head group (Dickson and Hille, 2019; Falkenburger et al., 2010). By contrast, the significance of the attached fatty acids has remained unappreciated (Barneda et al., 2019; Traynor-Kaplan et al., 2017). In mammalian cells, approximately 40-80% of the PIP molecules contain stearoyl/arachidonyl in the *sn*-1 and *sn*-2 positions, respectively designated as C18:0/C20:4 or 38:4 for the complete molecule (Falkenburger et al., 2010). A previous study revealed that mutations in the *p53* gene expand PIs containing short-length acyl chains, corresponding to 36-carbon PIs (Naguib et al., 2015). Remarkably, the shift in PI-lipids was not reflected in the lipid spectra of phosphatidylcholine (PC), asserting that the modification is not distinctive of all phospholipids (Naguib et al., 2015). Notably, both PC and PI share a common precursor, phosphatidic acid (PA). However, the latter steps in the pathway involve distinct enzymes to generate either PI or PC. Our studies revealed that *Abcb1*-deficient cells expressed comparable levels of PI synthase as well as the enzymes involved in PI acyl chain remodelling thus excluding these as a probable reason. As a result, it remains unclear as to how *Abcb1*-deficient cells acquire a distinct composition of PI-lipid chains. It is tempting to speculate that the precursor PA possessing a different lipid content gets accumulated in distinct membranes which is spatially only accessible to PI enzymes in deficient cells. Alternatively, it is also possible that the mature PIs are able to remodel their lipid chains once they have formed. If true, it will be interesting to decipher the mechanisms that prompt this variation.

Our study suggests that ABCB1 is critical in retaining immune equilibrium and the loss of ABCB1 dampens inflammasome activation. Keeping in view that aberrant PI signalling (including by the PI3K/AKT pathway) is associated with malignancies, it is plausible that ABCB1 does not exclusively regulates PI-lipid profile, and there exists redundancy in this pathway acting as a fail-safe mechanism. Accordingly, supplementation with linoleic acid similarly altered the PI profile incorporating elevated arachidonic acid levels. Notably, this phenomenon did not occur in cells directly exposed to arachidonic acid. Arachidonic acid has been shown to exhibit potent pro-apoptotic effects on macrophages by causing cell cycle arrest (Shen et al., 2018). In agreement, we observed a limited range in which macrophages tolerated arachidonic acid (data not shown). Linoleic acid is the most highly consumed polyunsaturated fatty acid found in the human diet. It is also the parent compound for the family of ω6 polyunsaturated fatty acids including arachidonic acid, which is further converted to a range of bioactive compounds such as leukotrienes, prostaglandins, and eicosanoids. Together, this suggests that linoleic acid is the preferred pathway for arachidonic acid expansion at the PI *sn*-2 position.

The activation of effector responses upon TLR ligation relies on signalling cascades involving TIRAP/MyD88 or TRAM/TRIF adaptor molecules which activate NF-kB or IFN-dependent signalling. Our data revealed that modified PI lipid chains elicit rapid degradation of the adaptor protein TIRAP. Bound to PI(4,5)P2 -enriched plasma membrane regions or PI(3)P at the endosomes, TIRAP surveys these compartments for activated TLRs. Notably, interaction with PIPs at its N-terminal phosphoinositide-binding domain is critical for TIRAP membrane recruitment and retention. A recent study demonstrated the participation of basic and nonpolar residues in the TIRAP phosphoinositide-binding domain. Significantly, the authors found both the inositol head group and acyl chains as critical in binding to TIRAP (Zhao et al., 2017). Under conditions of reduced phosphoinositide binding, IRAK1/4 phosphorylated Thr28 residue within the phosphoinositide-binding motif leading to TIRAP ubiquitination and degradation (Zhao et al., 2017). Independently, another study demonstrated that the ratio of PIP2 to PIP3 at the plasma membrane influenced TLR signalling (Aksoy et al., 2012). A decrease in PIP2 abundance at the plasma membrane concurrently resulted in TLR4 internalization and TIRAP redistribution to cytoplasmic compartments where it was degraded by the proteasome and calpain. Altogether, it is tempting to speculate that the modified PI is either incompetent in TIRAP binding and or fails to localize to the plasma membrane resulting in TIRAP ubiquitination and degradation.

NLRP3 inflammasome is regulated both at the transcriptional and post-translational levels. In particular, NLRP3 phosphorylation at distinct sites may either activate or inhibit the inflammasome. We observed that the increased incorporation of long-chain arachidonic acid by PI in WT cells through linoleic acid supplementation resulted in Ser293 phosphorylation in the NACHT domain and dampened inflammasome activation. Previous studies have shown increased phosphorylation at this site (Ser295) in THP-1 cells as a result of protein kinase A activation by prostaglandin E2 (Mortimer et al., 2016). Considering that linoleic and arachidonic acid are both precursors to prostaglandins, it may be interesting to test the overall basal levels of prostaglandins in these cells. Nevertheless, our data reveal the regulation of NLRP3 inflammasome by PI synthesis and metabolism at both the priming and activation steps.

Immune signalling triggers an adjusted inflammatory response, and any overt activation of these pathways may result in collateral damage. Therefore, it needs to be calibrated by mechanisms that offer a variable degree of responses and are infallible. While the phosphorylation of the inositol head group may at most result in seven distinct PIP species, the possibility to add an array of fatty acids, with different carbon lengths and saturation status, at the two positions on the PI glycerol linker offers far greater flexibility in terms of functions the cellular PIs can serve. In conclusion, our study provides insights as to how changes in PI lipid profile modify inflammasome activity and advances our understanding of the crosstalk between lipid metabolism and immune signalling.

## Acknowledgements

This work was supported in part by grants from The Wellcome Trust (108248/Z/15/Z), The Medical Research Council, UK (MR/S00968X/1) and core funds from Imperial College London to P.K.A. The authors declare no competing financial interests.

## Author Contributions

C.H., A.O., S.C., G.L-M. and P.K.A. designed experiments; C.H., A.O., S.L., and K.M-R. performed experiments; C.H., A.O., S.C., G.L-M. and P.K.A analyzed data; P.K.A. and G.L-M. provided resources; P.K.A. wrote the original draft, and all authors approved the final version. P.K.A provided overall supervision.

## Materials and Methods

### Ethics Statement

Experiments involving animals were performed in accordance with the Animals (Scientific Procedures) Act 1986, in accordance with a current UK Home Office licence and with approval from the Imperial College Animal Welfare and Ethical Review Body (AWERB).

### Bone-marrow derived macrophage isolation and cell culture

Bone marrow obtained from C57BL/6 mice (Charles River, UK) was isolated from the femurs and tibias of 6-8 week old mice as previously described (Anand et al., 2012, 2011). Bone marrow cells were then incubated in Dulbecco’s Modified Eagle Medium (DMEM) containing 10% heat-inactivated fetal bovine serum (FBS), 1% penicillin/streptomycin, 1% HEPES, and 30% conditioned media from L929 fibroblasts for 5 to 6 days at 37 °C and 5% CO_2_ for macrophage differentiation. Wild-type (WT) immortalised BMDMs (iBMDMs) were kindly provided by Katherine Fitzgerald and grown in DMEM containing 10% FBS, 1% penicillin/streptomycin, 1% HEPES, and 10% L929 conditioned medium. Cells were incubated and grown at 37 °C, 5% CO_2_ and passaged every 2-3 days. HEK293T cells were grown in DMEM containing % FBS and 1% HEPES at 37 °C, 5% CO_2_. Cells were grown until confluent and passaged every 2-3 days.

### Generation of CRISPR-Cas9 ABCB1b knock-out cells

Guides targeted exon 10 and exon 11 of the *Abcb1b* gene were designed using the CHOPCHOP online software (http://chopchop.cbu.uib.no/) and the Zhang Lab CRISPR design software (http://crispr.mit.edu/). The guides contained a *BsmBI* overhang and were as follows: Exon 10 gRNA – CACCG<u>AAGCCTTTGCAAACGCACGA</u> and its reverse complement <u>AAACTCGTGCGTTTGCAAAG</u>GCTTC and exon 11 gRNA – CACCG<u>CCCATCGAGAAGCGAAGTTC</u> and its reverse complement – <u>AAACGAACTTCGCTTCTCGA</u>TGGGC. Guide RNAs were annealed and ligated into the LentiCRISPRv2 plasmid (Addgene #52961) at the *BsmBI* restriction site. The recombinant plasmids (LentiCRISPRv2.gRNA10, LentiCRISPRv2.gRNA11) were then transformed into competent *Escherichia coli* (NEB) according to the manufacturer’s instructions. 1.85 μg of LentiCRISPRv2gRNA10 plasmid or LentiCRISPRv2gRNA10 plasmid, 0.42ug pVSVg plasmid and 1.3 μg psPAX2 plasmid were together transfected into the HEK923T cells using polyethyleneimine (PEI; Sigma), at a ratio of 1 μg DNA: 3 μg PEI. Supernatants containing lentiviral particles from transfected HEK293T cells were harvested at 48 hours post-transfection. 300 μl of lentiviral particle-containing supernatant was then added to the iBMDMs and cells were incubated at 37 °C for approximately 16 hours after which fresh media was added to cells. Forty-eight hours following transduction, puromycin (6 μg/ml, Sigma) was added for 7-10 days to select cells that had been successfully transduced. Puromycin-resistant cells were then trypsinised and seeded into a 96-well plate at an approximate concentration of 1 cell per well to obtain a clonal cell population. Single-cell colonies were identified and expanded followed by Sanger sequencing to identify mutations in Exon 10 or 11 of the *Abcb1b* gene. 2 clones, *Abcb1b*^-/-^ #1 and #2 were identified with deletions of 32 and 9 base pairs in exon 11, respectively (**Fig S2**).

### Cell stimulations

Primary and immortalised macrophages were seeded into either 6-well plates at a concentration of 2.5 × 10^6^, 12 well plates at a concentration of 1 × 10^6^ or 24-well plates at a concentration of 0.5 × 10^6^ cells per well. Where indicated, macrophages were treated with elacridar (1-10 μM, SML0486 Sigma) or Ko143 (50–200 ng/ml, K2144, Sigma) for 16 hours. In experiments with MβCD (C4555, Sigma), MβCD (5 – 10 μM) was added 30 minutes before the addition of LPS. In experiments with MG132 (10 μM), it was added 30 minutes before the addition of LPS. For fatty acid supplementation, distinct fatty acids were conjugated to albumin as previously described (Hunt et al., 2004). Cells were continuously cultured with albumin-bound arachidonic acid (5 μM), linoleic acid (20 μM), or a combination of arachidonic acid (5 μM) and stearic acid (20 μM) for at least two weeks prior to mass-spec analysis or any further experiments.

### Inflammasome activation

To activate the NLRP3 inflammasome, cells were incubated with LPS (500 ng/ml, Invivogen) for 4-6 hours to prime the macrophages, followed by ATP (0.5 μM, Sigma) or Nigericin (20 nM, Tocris) for approximately 45 minutes. For NLRC4 inflammasome activation, the *Salmonella Typhimurium* strain SL1344 was cultured overnight in 5 ml Luria-Bertani (LB) broth at 37 °C and on a shaker at 220 rpm. The bacteria were then added to the indicated cells, treated with or without elacridar (1 – 5 μM) or *Abcb1b*^-/-^ cells, at an MOI of 2 and incubated for approximately 4 hours. For AIM2 inflammasome activation, macrophages were transfected with 1 μg of poly(dA:dT) (Invivogen) complexed with Lipofectamine 2000 at a 1:3 ratio according to the manufacturer’s instructions for approximately 4-5 hours.

### Cell signalling experiments

The indicated cells were seeded into 6-well plates at a concentration of 2 × 10^6^ cells per well and incubated with elacridar (5 μM) overnight. The following morning media was replaced and cells were stimulated with LPS (500 ng/ml) for the following time points: 0 (no LPS control), 0.5, 1, 2 and 4 hours. Supernatants were discarded from each well and the cells were subsequently washed with PBS and cells were collected in RIPA buffer. Samples were incubated on ice for approximately 30 minutes before being centrifuged at 15000X g for 15 minutes at 4°C to remove nuclei. The supernatant was collected, and the protein concentration of each sample was measured using the BCA Protein Assay kit (Thermo Scientific #23227) according to the manufacturer’s instructions. Samples were all standardised to 1 μg/μl before immunoblot analysis.

### Immunoblot analysis

For immunoblot of phosphospecific antibodies, cells were collected in RIPA lysis buffer containing both protease and phosphatase inhibitors (Roche) and standardised to 1 μg/μl as mentioned above. Samples were boiled at 95 °C for 5 minutes before being resolved on 12% SDS-PAGE gels. For immunoblotting of caspase-1, NLRP3, IL-1β, ASC, GSDMD and GAPDH, lysates were collected in cell lysis buffer containing NP-40, DTT, and protease inhibitors. Samples were boiled at 95 °C for approximately 20-30 minutes before being resolved on 12% SDS-PAGE gels. SDS-PAGE gels were transferred to nitrocellulose membranes (GE Life Sciences) and subsequently blocked in a 5% milk solution in TBS-Tween (0.05%). Membranes were then incubated with the primary antibody overnight at 4 °C followed by incubation with the HRP-conjugated secondary antibody at room temperature for 1 hour. The primary antibodies used were as follows: P-glycoprotein (1:1000, Invitrogen MA1-2652), Caspase-1 (1:2000, AdipoGen #AG-20B-0042-C100), NLRP3 (1:2000, AdipoGen #AG-20B-0014-C100), ASC (1:2000, Adipogen #AG-25B-0006, AL177), GSDMD (1:1000, Abcam #ab209845), IL-1β (1:500, Cell Signalling #12426), GAPDH (1:2500, ThermoFisher # MA5-15738), Ikβα (1:1000, Cell Signalling, #9242), phosphor-IKβα (1:1000, Cell Signalling, #2859S), p38 MAPK (1:1000, Cell Signalling, #9212), phospho-p38 MAKP (1:1000, Cell Signalling, #4511), AKT (1:1,000; 4691; Cell Signaling Technology), phospho–AKT (1:1,000; 4060; Cell Signaling Technology), p70-S6K1 (1:1,000; 2708; Cell Signaling Technology), phospho-p70-S6K1 (1:1,000; 9234; Cell Signaling Technology), TIRAP (1:1000; Thermofisher; #PA5-88657), phospho-NLRP3 (1:1000, Thermofisher, # PA5-105071). The HRP-conjugated secondary antibodies (ThermoFisher) were used at 1:5000. Following secondary antibody incubation, proteins were visualised using either BioRad Clarity ECL substrate (BioRad #1705060) or the Pierce ECL Western Blotting Substrate (Thermo Scientific #32209) and processed on a Bio-Rad imager. Images were obtained using the Bio-Rad software, ImageJ.

### ELISA

Cell culture supernatants were measured for IL-1β (eBioscience #88-7013-88), TNF-α (eBioscience #88-7324-88) and IL-18 (MBL #7625), using ELISA kits according to the manufacturer’s instructions.

### Real-time PCR

RNA was isolated using TRIzol (Sigma # T9424) according to the manufacturer’s instructions. 250 μg of RNA from each sample was then reverse transcribed into cDNA using the High-capacity cDNA Reverse Transcription kit (Applied Biosystems #4368814) according to manufacturer’s instructions. Real-time PCR was then performed using the specific primers detail in **Table 1**. Real-time PCR was performed on an ABI7500 or ABI7900HT (Applied Biosystems) fast real-time PCR instrument.

**Table 1.**
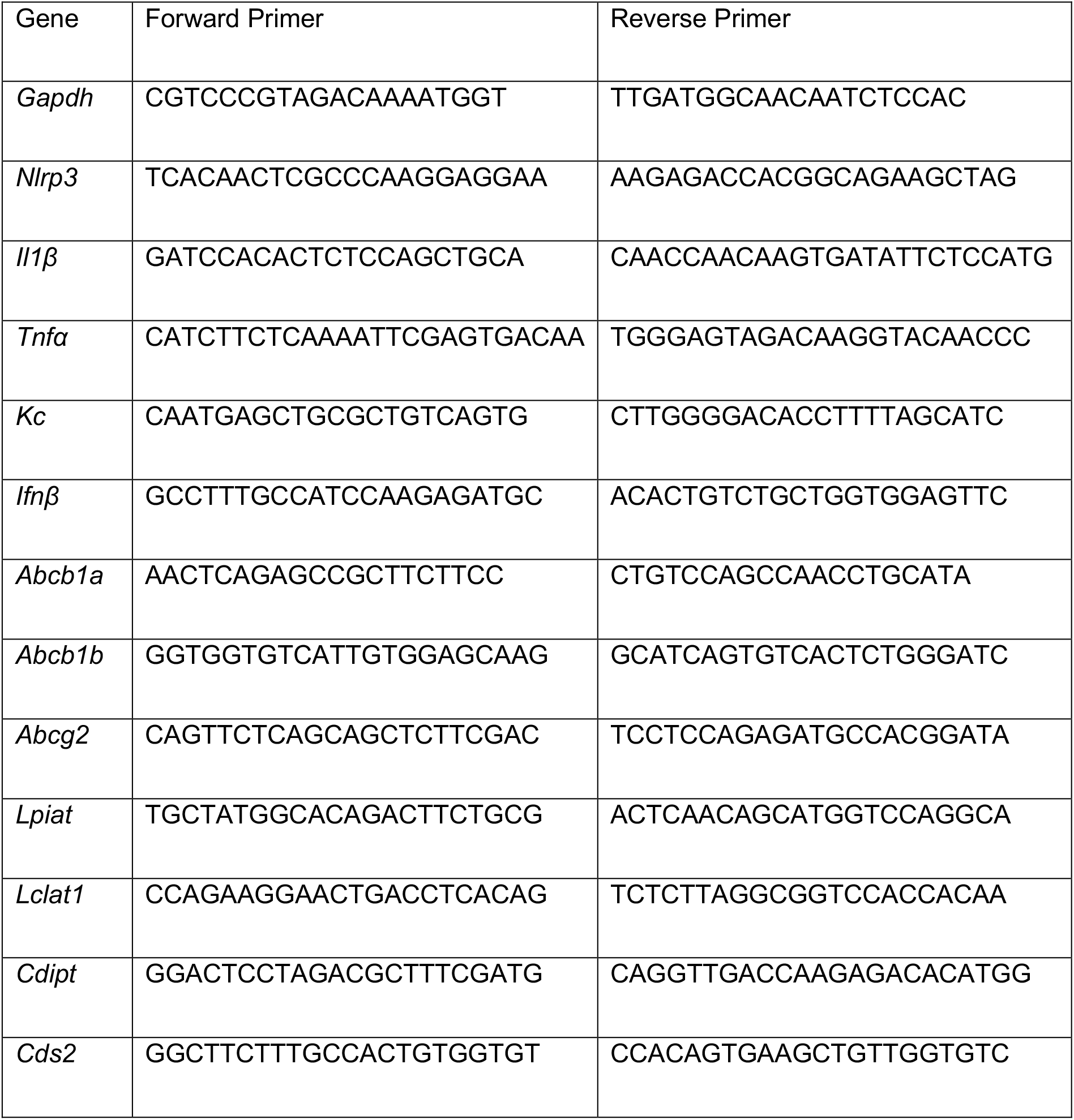

### Rhodamine 123 accumulation assay

BMDMs or *Abcb1b*^-/-^ clones were seeded into 96-well plates at a concentration of 1 × 10^5^ cells per well. Where indicated, BMDMs were treated with elacridar overnight (2-10 μM). Rho123 (2 μM, Sigma) was added and cells were incubated at 37 °C for 30 minutes. Cells were then washed with PBS 3 times to remove any extracellular Rho123 and DMEM media was replaced, after which cells were incubated for a further 30 minutes at 37 °C. Cells were then washed again with PBS and lysed in cell lysis buffer before being read at excitation and emission wavelengths of 485 and 535 nm, respectively in a fluorescent plate reader.

### Total cholesterol measurement

Total cholesterol was measured using the Cholesterol Amplex Assay kit according to the manufacturer’s recommendations.

### Filipin staining

WT and *Abcb1b*^-/-^ cells were seeded onto coverslips at a concentration of 4 × 10^5^ cells per well. After they had adhered, cells were fixed in 4% paraformaldehyde for 1 hour at room temperature. Cells were then stained with 25 μg/ml filipin (F9765, Sigma) overnight and washed 3 times with PBS before mounting on glass slides. Images were visualised on a Leica SP5 confocal microscope using 405 nm excitation and processed using the ImageJ programme.

### GM1 staining

Lipid rafts were assessed by specific labelling of endogenous GM1 ganglioside (a lipid raft marker) with the Vybrant Alexa Fluor 488 Lipid Raft labelling kit which makes use of fluorescently conjugated-CTB. The procedure was carried out according to the manufacturer’s instructions (Thermofisher). Samples were imaged using a Leica SP5 confocal microscope. Flow cytometry was performed using the Attune NxT Flow Cytometer (Thermofisher) and mean fluorescence intensity (MFI) was calculated using FlowJo™ v10.

### Caspase-1 inflammasome assay

In some experiments, caspase-1 activity in the cell culture supernatants was measured using Caspase-Glo® 1 inflammasome assay according to the manufacturer’s instructions (Promega). Luminescence was measured using an Omega plate reader.

### Immunofluorescence

WT, *Abcb1b*^-/-^ and linoleic acid -supplemented cells were seeded onto coverslips at a concentration of 4 × 10^5^ cells/well. After the experimental conditions, cells were fixed in 4% paraformaldehyde for 1 hour at room temperature and washed twice with PBS-glycine (50mM). The coverslips were blocked by incubating them for 20 minutes with PBS containing 1% BSA. Cells were then labelled with either anti-ASC antibody (1:100; AG-25B-0006, AL177; AdipoGen) or anti-TIRAP antibody (1:100; Thermofisher; #PA5-88657) for 1h at room temperature. Coverslips were then washed with PBS twice before incubating them with secondary antibodies (Thermofisher). Actin staining was carried out by labelling samples with Alexa Fluor™ 647 -conjugated phalloidin (Thermofisher; A22287) for 30 minutes. Coverslips were washed 3 times and mounted on slides using Fluoromount-G™ mounting medium with DAPI (Thermofisher; 00-4959-52). Images were acquired on an SP5 confocal microscope and analysed using ImageJ software.

### MALDI-MS Analysis

WT and *Abcb1b*^-/-^ macrophages were cultured in DMEM before being trypsinised and analysed via mass spectrometry. Prior to analysis, the super-2,5-dihydroxybenzoic acid (Sigma-Aldrich, catalogue no. 50862) matrix was added at a concentration of 10 mg/mL in a chloroform/methanol mixture at a 90:10 (v/v) ratio; 0.4 μL of a cell solution at a concentration of 2 × 10^5^ to 2 × 10^6^ mL^−1^, preliminary washed three times with double distilled water. corresponding to ~100–1000 cells per well of the MALDI target plate (384 Opti-TOF 123 mm × 84 mm AB Sciex NC0318050, 1016629), and 0.8 μL of the matrix solution were deposited on the MALDI target plate. These were then gently mixed with a micropipette, and left to dry. MALDI-TOF MS analyses were performed on a 4800 Proteomics Analyzer (with TOF-TOF Optics, Applied Biosystems) using the reflectron mode. Samples were analyzed in the negative ion mode operating at 20 kV. A total of three independent experiments were performed. Data obtained from mass-spec were analyzed using Data Explorer version 4.9 (Applied Biosystems), and assignments were based on the MS/MS fragmentation profile.

### Statistical Analysis

GraphPad Prism 9.0 software was used for data analysis. Data are represented as mean ± SD and are representative of experiments done at least three times. Statistical significance was determined by unpaired Student’s *t-test; p*<0.05 was considered statistically significant.

