## Supplementary Figure Legends for "NLRP3 inflammasome priming and activation are regulated by a novel phosphatidylinositol-dependent mechanism"

**S1 Fig. ABCB1 gene expression in mouse macrophages.** cDNA from control and LPS-primed (4 hours) immortalised BMDMs was amplified using primers specific to **(A)** *Abcb1a* or **(B)** *Abcb1b*. PCR products were separated by gel electrophoresis. **(C)** Relative fold change in the mRNA expression of *Abcb1a* and *Abcb1b* mRNA expression in iBMDMs by real-time PCR. mRNA expression is shown relative to *Gapdh*. **(D)** Primary mouse BMDMs were analysed for *Abcb1b* gene expression by PCR and separated by gel electrophoresis. **(E)** WT iBMDMs were stimulated with or without LPS (500 ng/ml) for the indicated time points and analysed by real-time PCR for *Abcb1b* mRNA expression. **(F)** Lentiviral particles containing sgRNA against *Abcb1b* exons 10 and 11 were generated and transduced into iBMDMs. Single-cell colonies were then generated, and clonal cell lines were assessed for deletions in exon 10 and 11 of the *Abcb1b* gene by Sanger sequencing. Two clonal lines with mutations were identified (*Abcb1b*$^{-/-}$ #1 and #2). **(G)** *Abcb1b* mRNA expression from CRISPR/Cas9 *Abcb1b*$^{-/-}$ macrophages generated in A. Data shown are mean ± SD and is representative of at least three independent experiments. *, p=<0.05, ** p = <0.01, by Student's t-test.

**S2 Fig. ABCB1 regulates activation of the NLRP3 inflammasome. (A)** WT BMDMs were treated with increasing amounts of elacridar overnight (1, 2, 5, and 10 µM), followed by LPS priming (500 ng/ml) for 4 hours and ATP (5 mM) for approximately 45 minutes. Cell lysates were immunoblotted for caspase-1 and GAPDH. **(B)** Supernatants from cells treated as in A were analysed for IL-1β by ELISA. Data shown are mean ± SD and is representative of at least three independent experiments. **, p=<0.01, **** p = <0.0001, by Student's t-test.

**S3 Fig. ABCB1 affects transcriptional responses downstream of TLR2, TLR4, and TLR7 ligation.** WT and *Abcb1b*$^{-/-}$ #2 cells were stimulated with either LPS (500 ng/ml), Pam3 (1 µg/ml) or imiquimod (1 µg/ml) for 4 hours and analysed for mRNA expression of **(A)** *Il1β,* **(B)**

*Tnfα,* and **(C)** *Cxcl1 (KC)*. mRNA expression shown is relative to *Gapdh*. Data shown are mean ± SD, and experiments shown are representative of 3 independent experiments. n.s, not significant, *, p = <0.05; **, p = <0.01; ***, p = <0.001; ****, p = <0.0001, by Student's t test.

**S4 Fig. MS/MS mass spectra of peaks at *m/z* 861 (A) and *m/z* 885 (B).** The fragment at *m/z* 241 corresponds to the polar head group fragment assigned to as inositol phosphate minus water. The fragment at *m/z* 153 corresponds to glycerophosphate minus water. The fragment at *m/z* 281 represents the fatty acid anions of oleic acid. In spectra A, the fragment ion at *m/z* 579 indicate the neutral loss of 282 corresponding to oleic acid and the fragments ions at *m/z* 417 show an additional neutral loss of 162 corresponding to an inositol unit minus water from the lyso-PI fragment ion at *m/z* 579. In spectrum B, the fragment ion at *m/z* 579 indicates the neutral loss of 306 corresponding to an eicosadienoic acid and the fragment ion at *m/z* 417 shows an additional neutral loss of 162 corresponding to an inositol unit minus water from the lyso-PI fragment ion at *m/z* 579. The fragment at *m/z* 303 represents the fatty acid anion of eicosatetraenoic acid.

**S5 Fig. The relative levels of total iBMDM percent PI content in cells either untreated or exposed to indicated fatty acids.** Cells were either left untreated or exposed to arachidonic acid or linoleic acid for at least two weeks before whole-cell lipidomics was carried out. The ratio of a particular PI species to the whole-cell representation of all observed PI species was calculated and shown here in percentage.

**Fig S6. ABCB1 deficiency disrupts GM1-positive membrane microdomains. (A)** Percentage GM1 as a total of GM1 and GM2 peaks identified in Fig 4A. **(B)** WT and *Abcb1b*$^{-/-}$ cells grown on coverslips were stained with cholera toxin B (CTB) (1 μg/mL) for 10 minutes at 4 °C followed by incubation with Alexa Fluor 488-conjugated anti-CTB antibody for 15 minutes at 4 °C to reveal GM1 presence by confocal microscope. **(C)** WT cells either untreated or treated with MβCD (10 µm, 30 min) and *Abcb1b*$^{-/-}$ cells were stained with Cholera Toxin subunit B (CTB) as above. Fluorescence was analysed by flow cytometry and representative spectra are shown. **(D)** WT cells and *Abcb1b*$^{-/-}$ cells were stimulated with LPS (500 ng/ml) and stained with CTB as above. **(E)** MFI (Mean fluorescence intensity) quantification of cells treated as in C. **(F)** MFI (Mean fluorescence intensity) quantification of cells treated as in D. **(G)** WT iBMDMs were treated with MβCD (5 and 10 µM) for 30 minutes followed by treatment with LPS (500 ng/ml; 4 hours) and ATP (5 µM; 45 min). Cell lysates were collected and immunoblotted for NLRP3, ASC, and GAPDH. Supernatants from cells treated as in (G) were analysed for IL-1β **(H)** or TNF-α **(I)** by ELISA. **(J-L)** WT iBMDMs were treated with MβCD (10 µM) for 30 minutes followed by treatment with LPS (500 ng/ml) for 4 hours and analysed for mRNA expression of *Nlrp3*, *Il1b*, and *Tnfα.* Data shown are mean ± SD and is representative of at least three independent experiments. **, p=<0.01, *** p = <0.001, **** p = <0.0001, by Student's t-test.
