## Supplementary Figures for "NLRP3 inflammasome priming and activation are regulated by a novel phosphatidylinositol-dependent mechanism"

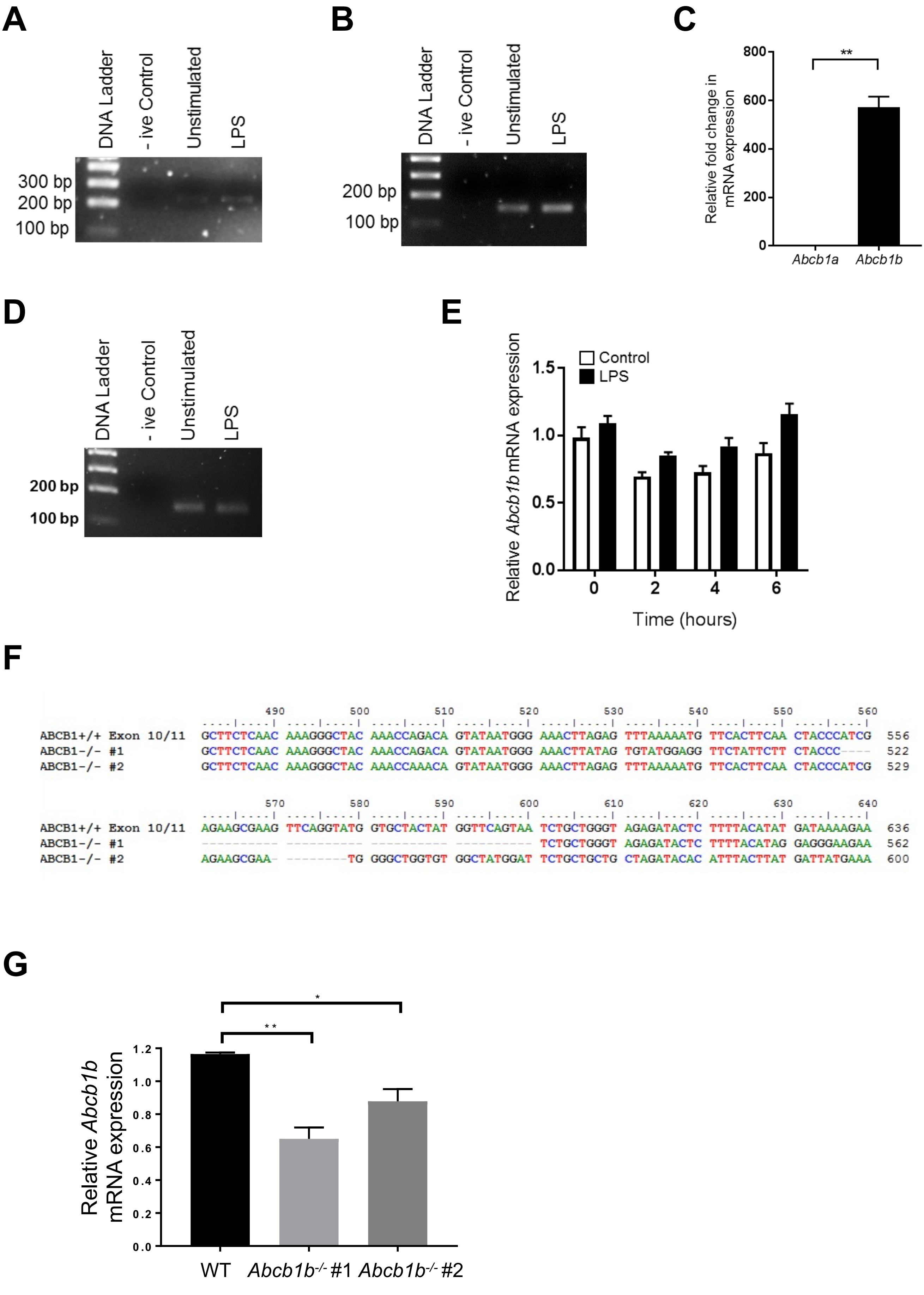
A
DNA Ladder
- ive Control
Unstimulated
LPS
300 bp
200 bp
100 bp
B
DNA Ladder
- ive Control
Unstimulated
LPS
200 bp
100 bp
C
Relative fold change in mRNA expression
800
600
400
200
0
**
Abcb1a
Abcb1b
D
DNA Ladder
- ive Control
Unstimulated
LPS
200 bp
100 bp
E
Control
LPS
Relative Abcb1b mRNA expression
1.5
1.0
0.5
0.0
0
2
4
6
Time (hours)
F
490 500 510 520 530 540 550 560
ABCB1+/+ Exon 10/11 GCTTCTCAAC AAAGGGCTAC AAACCAGACA GTATAATGGG AAACTTAGAG TTTAAAAATG TTCACTTCAA CTACCCATCG 556
ABCB1-/- #1 GCTTCTCAAC AAAGGGCTAC AAACCAGACA GTATAATGGG AAACTTATAG TGTATGGAGG TTCTATTCTT CTACCC---- 522
ABCB1-/- #2 GCTTCTCAAC AAAGGGCTAC AAACCAAACA GTATAATGGG AAACTTAGAG TTTAAAAATG TTCACTTCAA CTACCCATCG 529
570 580 590 600 610 620 630 640
ABCB1+/+ Exon 10/11 AGAAGCGAAG TTCAGGTATG GTGCTACTAT GGTTCAGTAA TCTGCTGGGT AGAGATACTC TTTTACATAT GATAAAAGAA 636
ABCB1-/- #1 ---------- ---------- ---------- ---------- TCTGCTGGGT AGAGATACTC TTTTACATAG GAGGGAAGAA 562
ABCB1-/- #2 AGAAGCGAA- --------TG GGGCTGGTGT GGCTATGGAT TCTGCTGCTG CTAGATACAC ATTTACTTAT GATTATGAAA 600
G
*
**
Relative Abcb1b mRNA expression
1.2
1.0
0.8
0.6
0.4
0.2
0.0
WT
Abcb1b-/- #1
Abcb1b-/- #2

### Figure S1

A

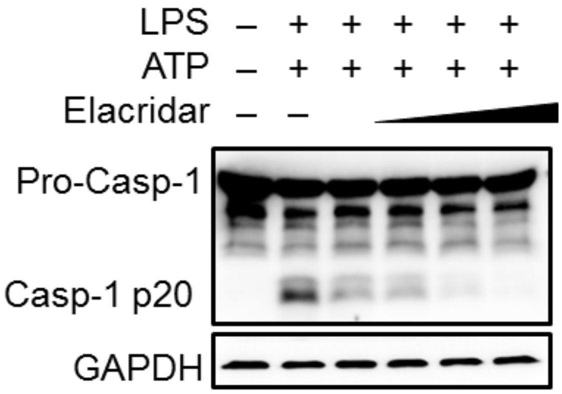
LPS – + + + + +
ATP – + + + + +
Elacridar – –
Pro-Casp-1
Casp-1 p20
GAPDH

B

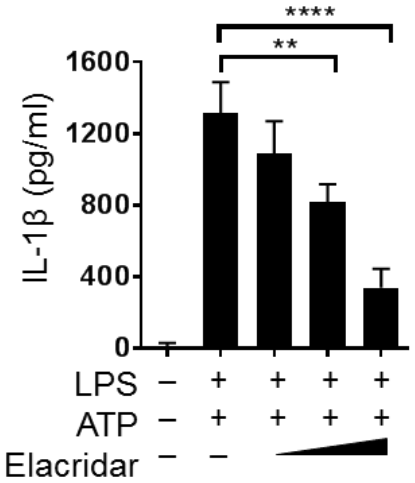
****
**
1600
1200
800
400
0
IL-1β (pg/ml)
LPS – + + + +
ATP – + + + +
Elacridar – –

**Figure S2**

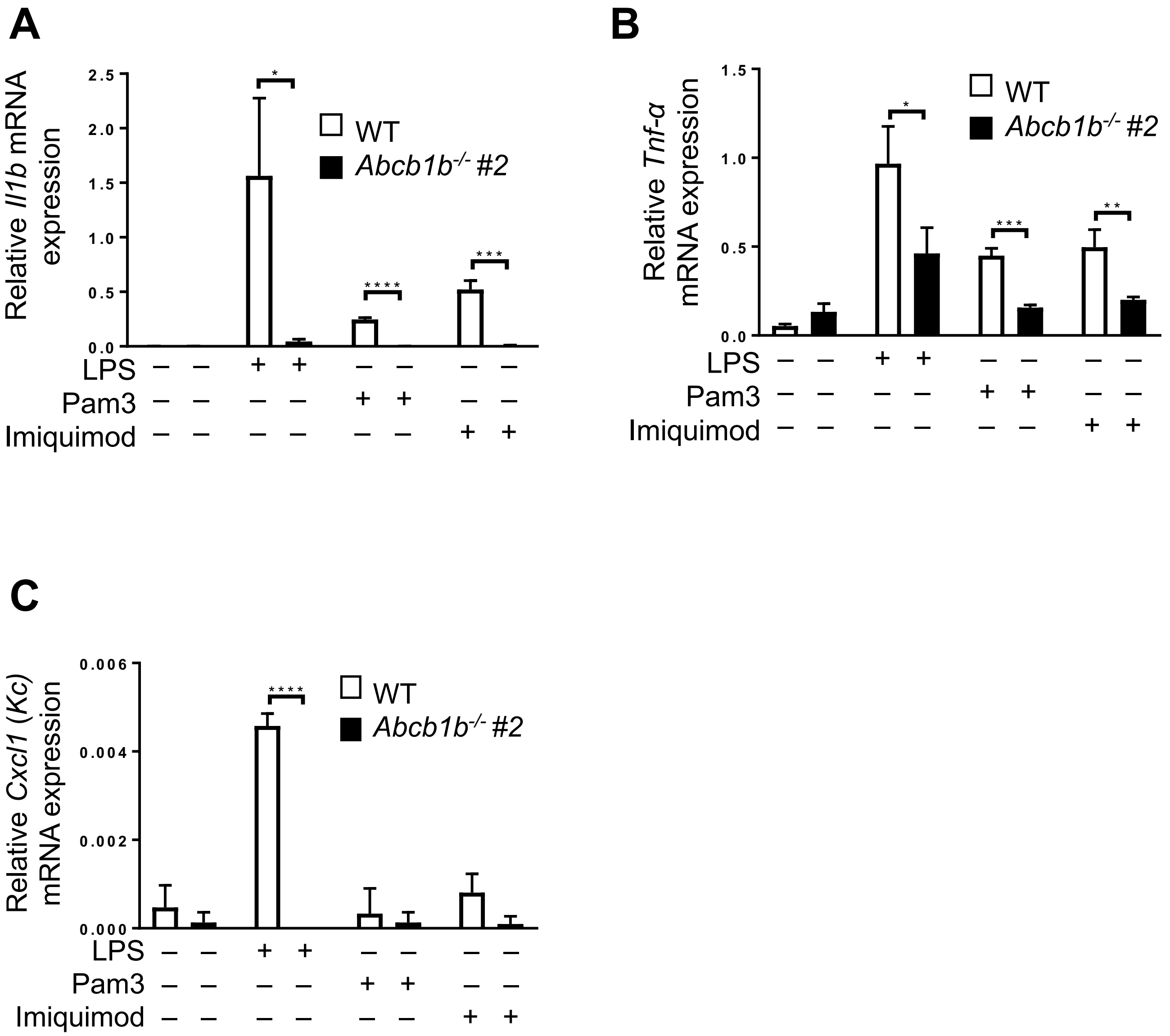
A
Relative Il1b mRNA expression
2.5
2.0
1.5
1.0
0.5
0.0
*
****
***
WT
Abcb1b-/- #2
LPS – – + + – – – –
Pam3 – – – – + + – –
Imiquimod – – – – – – + +
B
Relative Tnf-α mRNA expression
1.5
1.0
0.5
0.0
*
***
**
WT
Abcb1b-/- #2
LPS – – + + – – – –
Pam3 – – – – + + – –
Imiquimod – – – – – – + +
C
Relative Cxcl1 (Kc) mRNA expression
0.006
0.004
0.002
0.000
****
WT
Abcb1b-/- #2
LPS – – + + – – – –
Pam3 – – – – + + – –
Imiquimod – – – – – – + +

**Figure S3**

A

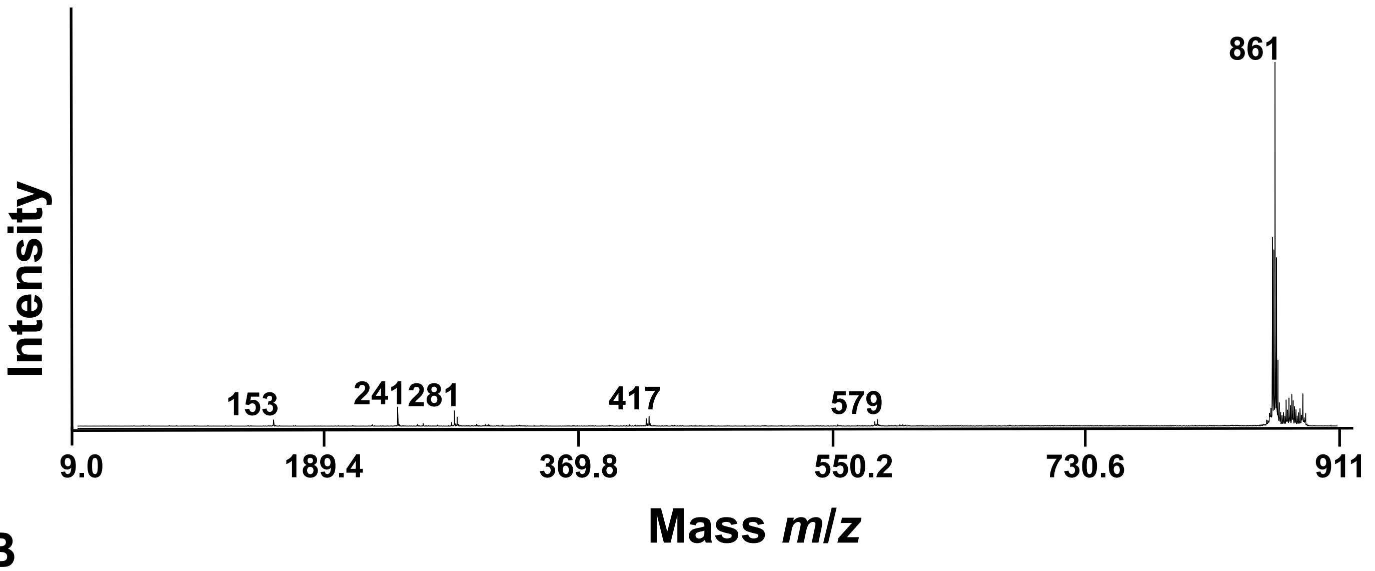
Intensity
861
153
241
281
417
579
9.0
189.4
369.8
550.2
730.6
911
Mass m/z

B

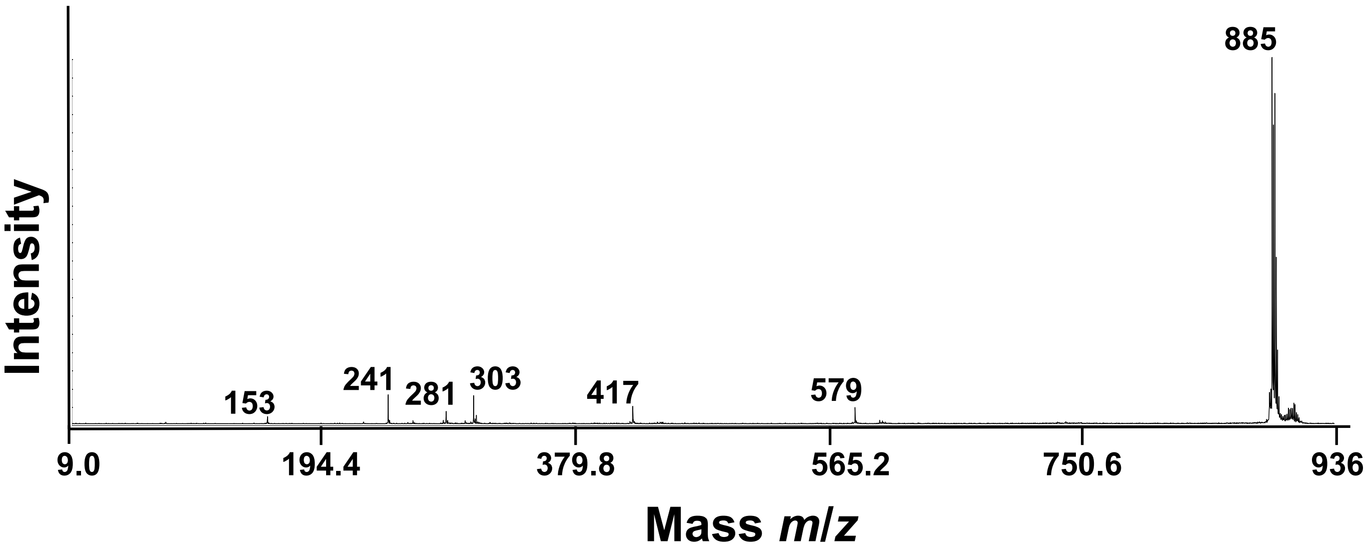
Intensity
885
153
241
281
303
417
579
9.0
194.4
379.8
565.2
750.6
936
Mass m/z

Figure S4

| Control | Mass (*m/z*) | | Dominant molecular species | Mean total iBMDM PI (%) |
|---|---|---|---|---|
| | 821 | 33:1 | | 5.57 |
| | 835 | 34:1 | 16:0/18:1 | 7.96 |
| | 861 | 36:2 | 18:1/18:1 | 39.75 |
| | 885 | 38:4 | 18:0/20:4 | 45.53 |
| **Arachdonic acid** | | | | |
| | 835 | 34:1 | 16:0/18:1 | 6.72 |
| | 861 | 36:2 | 18:1/18:1 | 38.29 |
| | 885 | 38:4 | 18:0/20:4 | 45.01 |
| **Linoleic acid** | | | | |
| | 835 | 34:1 | 16:0/18:1 | 3.26 |
| | 861 | 36:2 | 18:1/18:1 | 12.99 |
| | 885 | 38:4 | 18:0/20:4 | 69.01 |

**Figure S5**

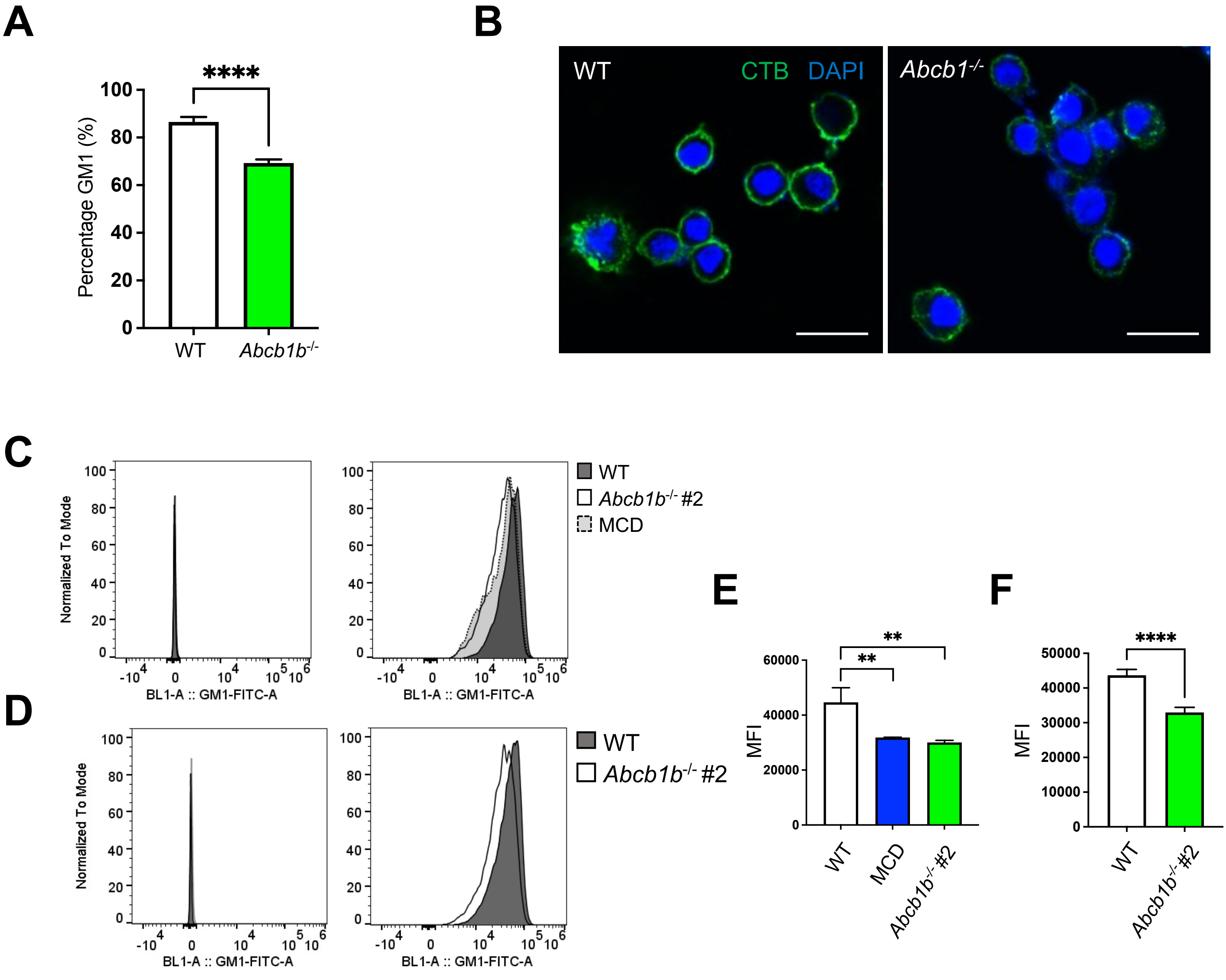
A
****
Percentage GM1 (%)
100
80
60
40
20
0
WT
Abcb1b-/-
B
WT
CTB DAPI
Abcb1-/-
C
Normalized To Mode
BL1-A :: GM1-FITC-A
WT
Abcb1b-/- #2
MCD
D
Normalized To Mode
BL1-A :: GM1-FITC-A
WT
Abcb1b-/- #2
E
**
**
MFI
60000
40000
20000
0
WT
MCD
Abcb1b-/- #2
F
****
MFI
50000
40000
30000
20000
10000
0
WT
Abcb1b-/- #2

**Figure S6**
